# Interpreting and de-noising genetically engineered barcodes in a DNA virus

**DOI:** 10.1101/2022.04.26.489490

**Authors:** Sylvain Blois, Benjamin M. Goetz, James J. Bull, Christopher S. Sullivan

## Abstract

The concept of a nucleic acid barcode applied to pathogen genomes is easy to grasp and the many possible uses are straightforward. But implementation may not be easy, especially when growing through multiple generations or assaying the pathogen long-term. The potential problems include: the barcode might alter fitness, the barcode may accumulate mutations, and construction of the marked pathogens may result in unintended barcodes that are not as designed. Here, we generate approximately 5000 randomized barcodes in the genome of the prototypic small DNA virus murine polyomavirus. We describe the challenges faced with interpreting the barcode sequences obtained from the library. Our Illumina NextSeq sequencing recalled much greater variation in barcode sequencing reads than the expected 5000 barcodes – necessarily stemming from the Illumina library processing and sequencing error. Using data from defined control virus genomes cloned into plasmid backbones we develop a vetted post-sequencing method to cluster the erroneous reads around the true virus genome barcodes. These findings may foreshadow problems with randomized barcodes in other microbial systems and provide a useful approach for future work utilizing nucleic acid barcoded pathogens.

## INTRODUCTION

The within-host population dynamics of a microbe are usually studied as population abundances across time and tissues (1,2). Although informative, this approach is blind to differences among individuals within populations. Thus, a virus concentration of 10,000 per mL maintained over time could be achieved if 10,000 lineages are all just maintaining themselves or could alternatively be achieved if 100 lineages are each producing 100 progeny and the other 9,900 are cleared by the host. Knowing those dynamics can shed light on the different processes that may be occurring during infection (e.g., latency of part of the population) and can reveal the extent of tissue subdivision of the infection and population bottlenecks. An understanding of dynamics may even inform treatment alternatives.

Technical advances in genetic engineering and sequencing now allow microbial populations to be established in which each individual microbe can be distinguished from nearly all others in an inoculum even though all individuals are the same genetic strain. With this technology, known as DNA barcoding, it becomes possible to create separate identities for potentially millions of individual lineages used to simultaneously infect a single host (3–10). Barcoding usually involves the insertion of short, randomized DNA segments in the genome, the number of different types increasing as 4^N^, where N is the number of bases in the randomized insert. Each lineage is then defined by its unique barcode, and all descendants from each infecting individual can be discriminated from descendants of other infecting individuals through inheritance of the barcode. In essence, barcoding enables a pedigree analysis of an ongoing population.

The analysis of barcodes is trivial when they do not change from parent to offspring. But barcodes may be prone to mutation and sequencing errors. In that case, analysis of ancestor-descendant relationships will require a statistical correction (clustering) to group non-identical barcodes that nonetheless share common ancestry. The suitability of any specific clustering algorithm may depend on the biological details of the implementation (11) – the various factors contributing to parent-offspring differences in barcode sequence as well as the extent of differences among the barcodes in the founding population. But the clustering principles should transcend specific applications.

Here, we describe the creation of barcoded polyomaviruses that were generated from a library of barcoded plasmids. When the sequences of the plasmid library were analysed, it became obvious that a substantial fraction of sequencing reads had to be errors. We applied a vetted barcode clustering algorithm to associate the erroneous barcode sequences with the true parent barcode. We utilized a series of defined experimental known barcode genome controls and simulations to empirically ascertain appropriate clustering parameters. This system afforded considerable control over and investigation of the factors that complicate the interpretation of barcode abundance with clustering proving essential to the interpretation of barcodes and their abundances. We characterize the virus library and the components that went into individual steps of its construction. The result is an in depth understanding of the functionality of different computational clustering parameters, linear dynamic range, and sensitivity of Illumina barcode quantification. These findings provide a functional approach applicable to experimental infections with murine polyomavirus (muPyV) and likely other barcoded microbes.

## MATERIAL AND METHODS

### Cells and viruses

#### Cells

The NMuMG cells (ATCC^®^, # CRL-1636) were kindly provided by Prof. Aron Lukacher and maintained in DMEM (Corning, # 10-013-CV) supplemented with 10% (v/v) fetal bovine serum (Corning, # 35-015-CV) and 1% (v/v) penicillin-streptomycin (Corning, # 30-002-CI) (12).

### Viruses

#### muPyV PTA wild-type stock generation

pBluescript-sk+ vector containing the full genome of the muPyV PTA strain (GenBank, accession No U27812) was digested by BamHI-HF (New England Biolabs, # R3136L) to separate the muPyV genome from the vector and gel purified. Then, the muPyV genome was self-ligated overnight at 16°C using the T4 DNA ligase (New England Biolabs, # M0202L) and purified prior transfection into NMuMG cells using the Lipofectamine™ 2000 reagent (Invitrogen, # 11668019). NMuMG cells were cultured until 90% cytopathic effect (CPE) was reached (5 days post-transfection). The cells and the supernatant were collected, subjected to freeze/thaw cycles and cleared by centrifugation at 4°C. Fresh NMuMG cells were infected with the crude lysate at 37°C and cultured until 90% CPE (day 6 post-infection). The infected cells and the supernatant were harvested, subjected to freeze/thaw cycles, buffered at pH 8.5, incubated at 42°C for 30 min, vortexed to release muPyV and cleared by centrifugation. The muPyV PTA stock was aliquoted and stored at -80°C until use.

#### Construction of the barcoded muPyVs library

pBluescript-sk+ vector containing the full genome of muPyV PTA (GenBank, accession No U27812) was amplified by reverse PCR using the Phusion^®^ High-Fidelity DNA Polymerase (New England Biolabs, # M0530L) and the 5’ phosphorylated primers BCFSB7 5’-NNNNNNCAATTGAATAAACTGTGTATTCAGCTATATTC-3’ and BCRSB7 5’-NNNNNNGAATAAACATTAATTTCCAGGAAATAC-3’ (Integrated DNA Technologies). The PCR product was digested by DpnI (New England Biolabs, # R0176L), gel purified and self-ligated using the T4 DNA ligase (New England Biolabs, # M0202L) before transforming MAX Efficiency^®^ DH5α™ competent cells (ThermoFisher, # 18258012). Transformed bacteria were cultured overnight in LB broth containing 100μg/mL of ampicillin and the plasmid library was purified. The number of transformed bacteria was estimated in parallel on an aliquot of the transformed bacteria by counting the colonies on LB agar plates containing ampicillin. The presence of the barcode was confirmed in plasmids from colonies by enzymatic digestion using BamHI-HF and MfeI-HF (New England Biolabs, # R3136L and # R3589L, respectively) and by Sanger sequencing using the primer FullSeq3 5’-GTTAGAGTGTATGATGGGACTG-3’. The virus library was then prepared by transfecting the plasmid library as described above for the generation of the muPyV wild-type stock. To confirm the presence of the barcode in the virus stock, virus DNA was purified using the QIAamp^®^ DNA Mini Kit (Qiagen, # 51304) and the region surrounding the poly A signals was PCR amplified using Taq DNA polymerase (New England Biolabs, # M0267L) and the primers NGS_Fwd 5′-CATGGCCTCCCTCATAAGTT-3′ and NGS_Rev 5′-GAATATAGCTGAATACACAGTTTATTC-3′ following the manufacturer’s recommendation. The PCR product was purified, cloned using the TOPO^®^ TA Cloning kit for Sequencing (ThermoFisher, # 450071) and Sanger sequenced (Supplementary Figures S1 and S2).

#### muPyV titration by immunofluorescence assay

NMuMG cells were infected in duplicate with serial dilutions of the virus sample for 1 hour at 37°C. 30-40 hours p.i., cells were gently washed once and fixed/permeabilized with PBS containing 4% paraformaldehyde. Cells were washed three times, blocked with PBS containing 1% goat serum overnight at 4°C and incubated for 1 hour at room temperature with a rabbit anti-PyV VP1 antibody (a gift from Richard Consigi) diluted 1:50 in PBS. After three washes, the cells were stained with Alexa™ Fluor 488 goat anti-rabbit IgG (ThermoFisher, # A-11008). The number of stained cells per field was counted under an inverted fluorescence microscope (Leica), and infectious titers (IU/ml) were calculated.

### Illumina NextSeq library preparation

#### Enrichment PCR

This first step PCR amplifies a 360 bp fragment encompassing the barcoded region of muPyV genome using primers NGS_Fwd 5′-CATGGCCTCCCTCATAAGTT-3′ and ReVOUT1 5’-CAGGGTCTTGTGAAGGAGGT-3’ (primers binding site shown in Supplementary Figure S3). Briefly, the enrichment PCR reaction contained 4.77 × 10^5^ copy number of muPyV DNA, 0.5µM of the above primers, 200µM of each dNTP, 1U of Phusion® High-Fidelity DNA Polymerase (New England Biolabs, # M0530L) in 1X of Phusion^®^ HF reaction buffer. After an initial denaturation step for 1 min at 98°C, the amplification was performed by 22 cycles of 10 sec at 98°C (ramp 2°C/sec), 15 sec at 55°C (ramp 2°C/sec), 30 sec at 72°C (ramp 2°C/sec) followed by a final extension step for 5 min at 72°C. The DNA polymerase was inactivated at -20°C. The presence of a specific amplification product was checked on a 2% agarose gel.

#### Indexing PCR

This second step PCR amplifies a 75 bp fragment encompassing the barcode region from the enrichment PCR product, adding Illumina adapters and indexes to permit multiplexing, as well as spacers to increase the library complexity. The amplicons size generated ranges from 211 to 216bp. The sequence of the staggered indexing primers (Integrated DNA Technologies) is presented in Table 1 (primers binding site shown in supplementary Figure S3). Briefly, enrichment PCR products were digested with 20U of ExoI (New England Biolabs, M0293L) at 37°C for 30 minutes to remove residual primers and heat inactivated at 80°C for 20 min before proceeding to the indexing PCR. Then, two indexing PCR reactions were made for each sample containing 1µl of the enrichment PCR products, 0.3µM of indexing primers, 200µM of each dNTP, 1U of Phusion® High-Fidelity DNA Polymerase (New England Biolabs, # M0530L) in 1X of Phusion^®^ HF reaction buffer. After an initial denaturation step for 1 min at 98°C, the amplification was performed by 22 cycles of 10 sec at 98°C (ramp 2°C/sec), 15 sec at 54°C (ramp 2°C/sec), 30 sec at 72°C (ramp 2°C/sec) followed by a final extension step for 5 min at 72°C. The DNA polymerase was inactivated at -20°C. The presence of a specific amplification product was checked on a 2% agarose gel.

**Table 1:**
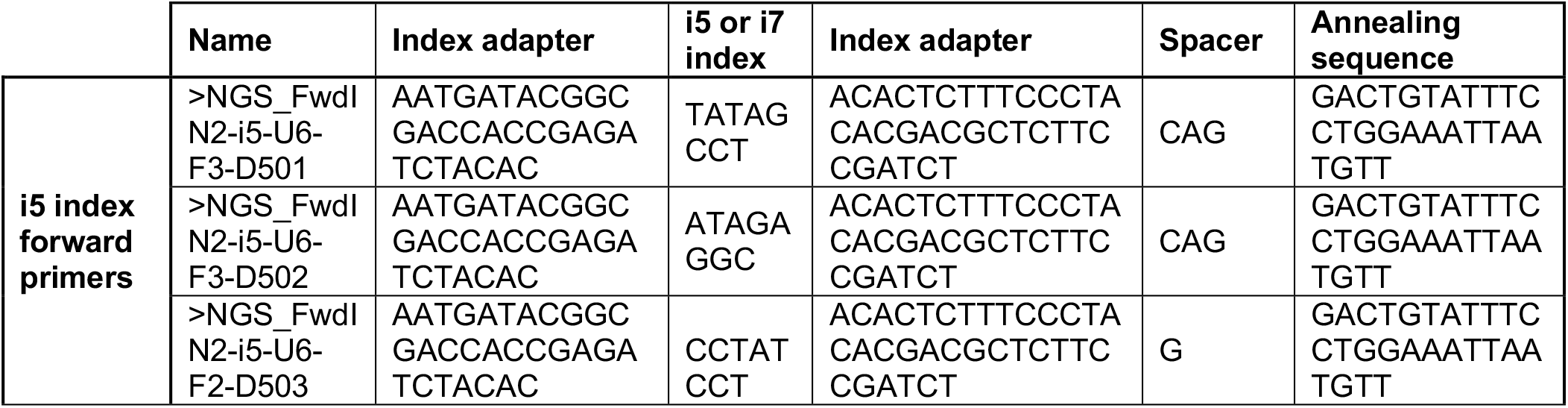
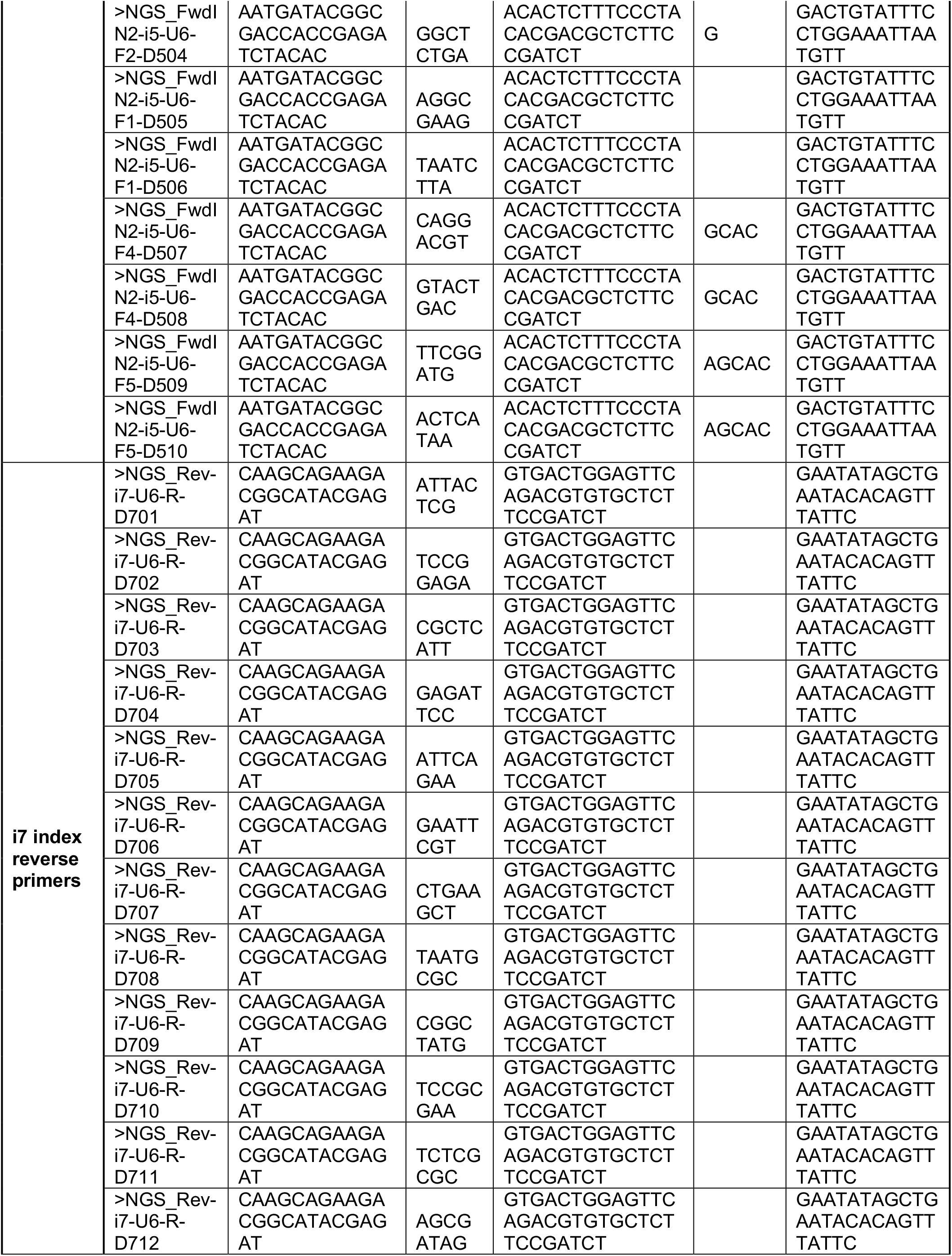
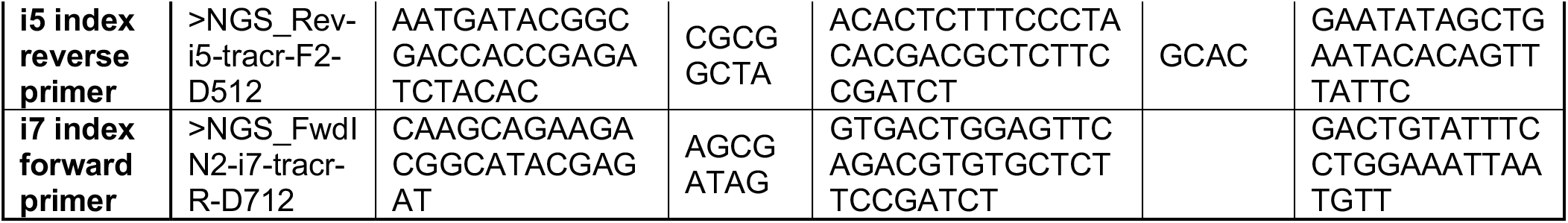
Staggered indexing primers.

### Amplicons pooling and quality control

The two indexing PCR reactions were merged together and concentrated using the Monarch^®^ PCR & DNA cleanup kit (New England Biolabs, # T1030L) before gel purification using the Monarch^®^ DNA gel extraction kit (New England Biolabs, # T1020L). The DNA concentration of each sample was determined using the QUBIT™ dsDNA BR assay kit (ThermoFisher, # Q32850). The samples were normalized and pooled together to constitute the final Illumina library. The final library was checked on a 2% agarose gel and sent to the Genomic Sequencing and Analysis Facility of the University of Texas at Austin for a Bioanalyzer (Agilent) quality control prior to the Illumina NextSeq SR75 sequencing run. To increase diversity, PhiX DNA was also included (∼5% to the first run and ∼34% to the second run).

### Extracting barcodes

Barcodes were extracted from FASTQ files off the sequencer using Cutadapt (13), with the default error allowance of 10% for the linked adapters (requiring both adapters to flank the barcode) (Figure 1).

**Figure 1.**
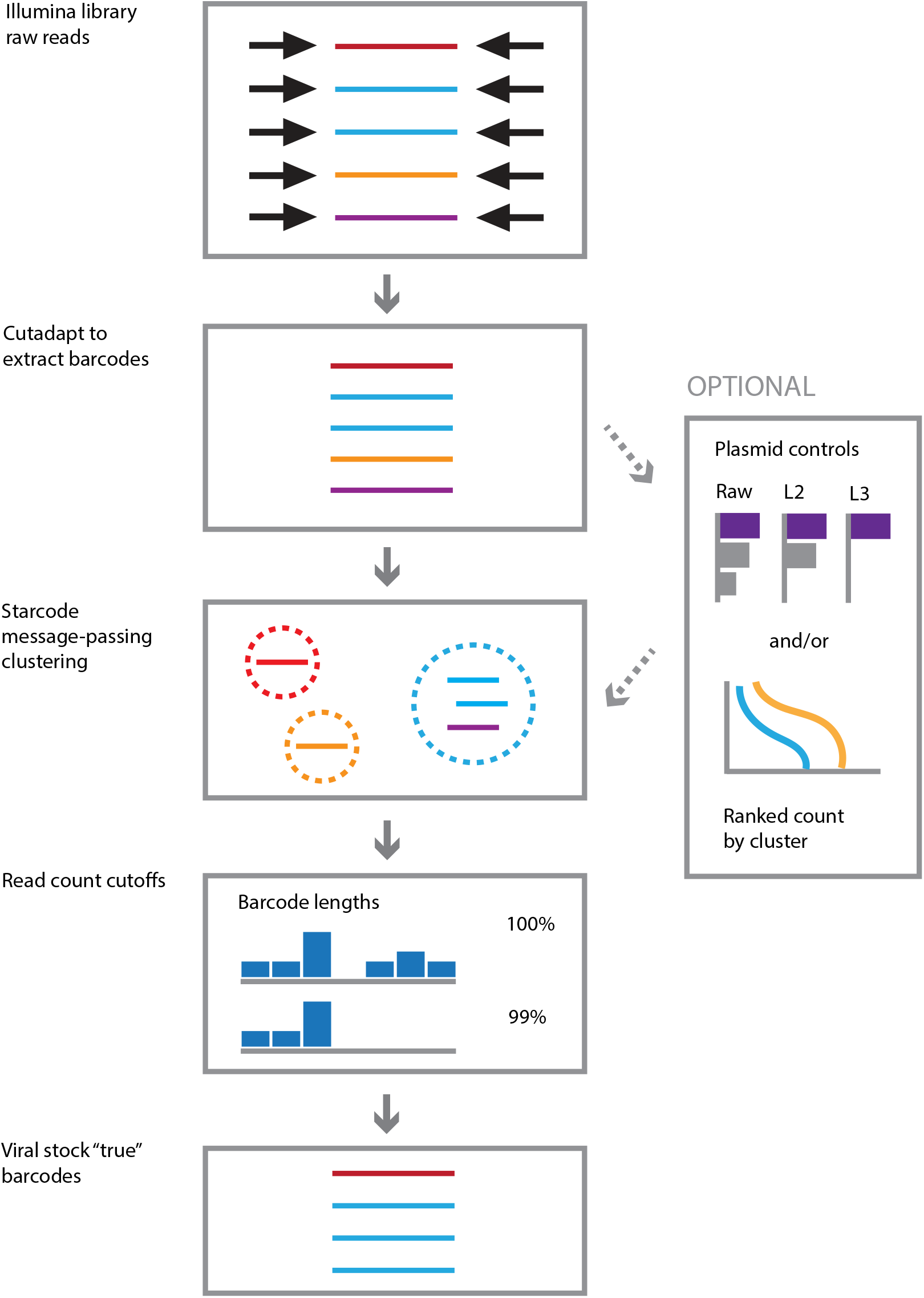
Outline of computational workflow for selecting stock barcodes. Linker sequences are trimmed with Cutadapt (colors represent distinct barcodes). Clustering is performed using the Starcode message-passing algorithm to determine centroids, defining “centroid” as the representative barcode (dotted circles represent clusters). We empirically determined which edit distance to employ by examining two criteria: 1) the performance of different edit distances on various amounts of plasmid controls, each containing a barcode of known sequence, to recall correct input barcode sequences and 2) “shoulder cutoff” using the plot line of the distribution abundance of called clusters from our barcoded plasmid library. In defining the “true” barcodes of our virus library, we utilized L=3 Levenshtein distance because it performed slightly better in our defined barcode plasmid controls over the default Starcode Levenshtein distance of L=2. Final “true” barcodes were then assigned by omitting the lowest abundance barcodes (omitting the bottom barcodes that account for a cumulative 1% of all counts), based on the assumption that the lowest abundance reads were most likely to be sequencing artifacts.

### Clustering barcodes

Unless otherwise noted, barcodes were clustered using Starcode (14) for message-passing clustering, using a Levenshtein distance of 3. The highest-count barcode in a cluster (the centroid) was used as the representative for the cluster (Figure 1).

### Checking theoretical overclustering via simulation

To check the possibility of choosing a Levenshtein distance of 3 leading to overclustering, we simulated 5000 random 12mers and calculated pairwise Levenshtein distances between them using the base R adist function.

### Selection of barcodes by cumulative count cutoff

The representatives of the clusters were cut off using a cumulative count criterion. Unless otherwise noted, the barcodes whose collective counts accounted for 99% of all counts were kept, and the rest discarded. Applying this threshold removed barcodes whose length was significantly more than 12 nt (Figure 1).

## RESULTS

### Generation of barcoded DNA viruses

In brief, we engineered the murine polyomavirus (muPyV) (dsDNA genomes of ∼5.4 kbp) to carry 18 bp inserts, 12 bases of which were randomized A, C, G, T with equal probability. We chose the non-protein-coding genomic region in between the opposing early and late polyadenylation sites for the insert, which was suggested as a viable location based on the tolerance of Bandicoot Papillomatosis Carcinomatosis Virus 1 (BPCV1) to a large natural genomic insert in an analogous region and a restriction site being successfully engineered in a similar location in a previously described murine PyV mutant (15–17). We cloned barcodes into bacterial plasmids containing the virus genome and then generated barcoded viruses as described in Methods. Barcoded virus stocks gave rise to high titers of virus, implying no overt fitness defects of these barcoded viruses (Table 2).

**Table 2:** Virus stocks titer (IU/mL) as determined by immunofluorescence assay.

|  | muPyV titer (IU/mL) |
| --- | --- |
| Wild-type virus | $7.14 \times 10^6$ |
| Barcoded virus library | $2.50 \times 10^6$ |

### An absolute bottleneck on barcode pool depth

We estimated the composition of the plasmid library that gave rise to the virus library to be composed of approximately 5700 total bacterial colonies, necessarily the maximum possible number of unique barcodes present. To get a sense of barcode similarity across different colonies, the barcodes of 21 separate colonies were Sanger-sequenced individually; 20 were unique. These findings rule out that one or a few barcodes dominate the population.

The barcode sequence repertoire that is represented in our plasmid library is sparse in the space of 4^12^ possibilities, with at least 2940 (4^12^ /5700) times more possible 12 mers than barcodes detected. As further addressed below, this sparseness allowed us to discover – and resolve – errors in the assaying of barcodes.

### Illumina sequencing exhibits elevated variation in barcodes

We utilized Illumina NextSeq SR75 sequencing of the virus population we generated from the plasmid library. Inspection of the Illumina reads of the barcode region showed 535,772 different sequences, far exceeding the number of plasmids (∼5700 bacterial colonies) used to generate the virus library. We observed that there were orders of magnitude variation in the abundances of the different barcodes. This variation must have arisen after the founding of the 5700 bacterial colonies. The steps where this discrepancy could arise include virus production, the Illumina library preparation, and Illumina sequencing itself. Therefore, the first question we addressed is whether Illumina sequencing generated excess barcodes in the plasmid library. Indeed, similar to the virus library, approximatively 274,213 different barcodes are detected in sequences of the plasmid library that gave rise to the virus library. This finding identifies procedural introduction of sequence variation in generating/analyzing barcoded virus libraries.

### Sequences of plasmid controls also reveal excess barcode variation

To resolve the magnitude and origins of barcode variation observed in library reads, we developed controls using plasmid DNA containing barcoded virus genomes. Plasmid DNA has the advantages that (i) DNA can be isolated from a single colony and thus is known to be limited to one barcode, (ii) a plasmid DNA can be directly sequenced with Sanger technology, such that the true, consensus sequence is known, and (iii) the DNA has not been manipulated with steps leading to virus creation and possible introduction of mutations.

The first controls used a single plasmid whose barcode region was sequenced with both Sanger and Illumina methods; alternative versions of this control varied the concentration of plasmid DNA and the number of enrichment PCR cycles used during the Illumina library preparation (Table 3). Illumina sequences from these single-plasmid controls revealed many barcode sequences (Figure 2), thus indicating that most of the barcodes identified were due to errors in and leading up to DNA sequencing. The percent of errors in raw data was high and ranged from 3.71 to 6.11% across different replicates (Table 4, top line of each plasmid entry). Altering the number of PCR cycles or input DNA concentration had little effect on the variation we observed.

**Table 3.** Single-plasmid controls preparation.

| Single-plasmid control | Barcode | Plasmid copies | PCR cycles |
| --- | --- | --- | --- |
| 1P-A | TCACAGGGGTAA | 1.00E+02 | 40 |
| 1P-B | TCACAGGGGTAA | 1.00E+03 | 35 |
| 1P-C | TCACAGGGGTAA | 1.00E+04 | 32 |
| 1P-D | TCACAGGGGTAA | 1.00E+05 | 29 |

**Figure 2.**
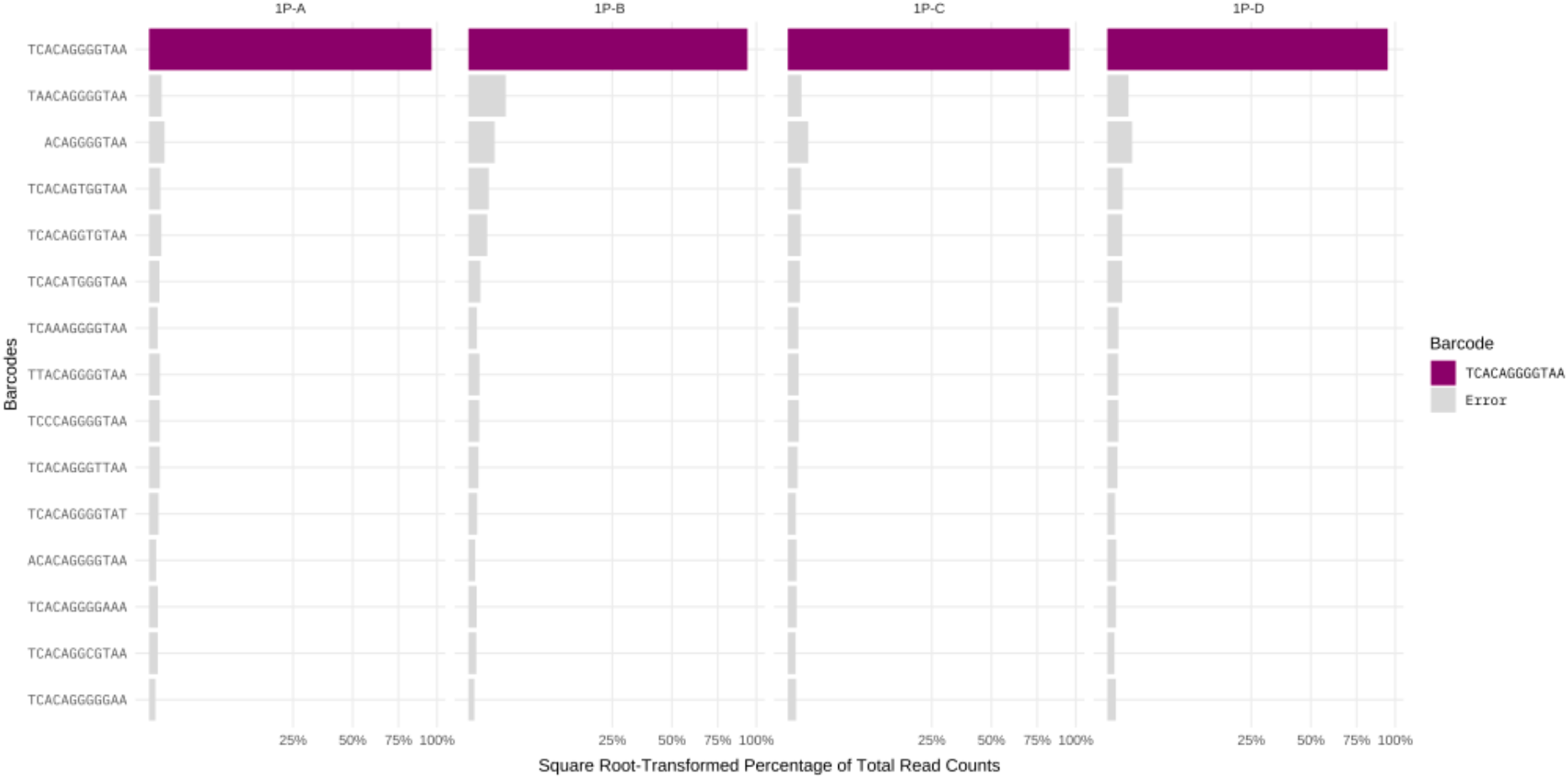
Illumina sequencing reads from single-plasmid controls. The control spiked-in barcode is highlighted in purple, erroneous barcodes are shown in gray. Shown are the top highest count 15 barcode sequences tallied across the four plasmid controls. Raw counts are shown with no clustering applied, with the x-axis square root-transformed to highlight low values.

**Table 4.** Percent of erroneous barcodes in Illumina sequencing reads of single-plasmid controls with or without clustering and different Levenshtein distances applied.

| Single-plasmid control | Clustering distances | Percent of erroneous barcodes |
| --- | --- | --- |
| **1P-A** | Raw | 3.71% |
|  | L1 | 0.51% |
|  | L2 | 0.06% |
|  | L3 | 0.03% |
| **1P-B** | Raw | 6.11% |
|  | L1 | 0.98% |
|  | L2 | 0.03% |
|  | L3 | 0.02% |
| **1P-C** | Raw | 4.23% |
|  | L1 | 0.73% |
|  | L2 | 0.07% |
|  | L3 | 0.04% |
| **1P-D** | Raw | 5.07% |
|  | L1 | 0.97% |
|  | L2 | 0.06% |
|  | L3 | 0.03% |

The high error rate is striking. Inspection of Figure 2 reveals the nature of deviations from the true barcode. For a 12-mer, there are approximately 37 possible 1-step deviations that ignore the identity of the change: 12 single-base changes, 13 single-base insertions and 12 single-base deletions. If the identity of the new base is accounted for, the 37 increases to 100 possible 1-step deviations. In these controls, we observed single-base changes and base deletions. If virus propagation was involved, this error rate could be explained as some type of virus intolerance of the insert. But this control was based on a plasmid containing a virus genome with a single barcode. Although we do not fully understand why we observe an apparent high error rate, we note that others assaying similarly homogenous amplicons also observe a higher error rate than is typical for Illumina (18). Thus, we believe it is likely that other barcode approaches with low complexity amplicons would encounter the same phenomena.

### Validation of clustering strategy, limit of detection, and linearity of defined plasmid controls

The goal of clustering is simply to correct the errors in barcode reads so that we know the true parents – as if there were no errors to begin with. This means we want the clustering method to group erroneous barcodes with their true sources. This task is easy with our control data because we know the source barcodes and thus know the true and erroneous ones. In contrast to controls, true barcodes are not necessarily known in experimental samples, thus, any clustering algorithm developed for the controls must be applicable to samples without known true barcodes.

As a sequence clustering algorithm, we utilized Starcode which is particularly suited to nearly identical length sequences, with a significant range of counts (14). For our initial implementation, we utilized Starcode’s message-passing algorithm (described in detail below). A crucial parameter for Starcode is the Levenshtein distance (L), which counts the number of simple ‘mutations’ (base changes, single base deletions, and single base insertions) required to render two sequences identical. For 12 nt sequences, Starcode’s default value for Levenshtein distance is L=2, although there is little published experimental evidence to justify which distance parameter is best for barcodes (see below). In order to test this, we used different Levenshtein distances on single-plasmid controls. Data reveals that L=1 eliminates 80%-90% of the erroneous barcodes. L=2 eliminates another 80-95%, and L=3 eliminates slightly more (Table 4).

We then developed a second set of controls using 10 plasmids containing known different barcoded virus genomes at different concentrations and we varied the number of enrichment PCR cycles used during the Illumina library preparation (Table 5 and supplementary Table S1). These controls take a step toward generating a typical dataset while still allowing detailed *a priori* knowledge of the source barcodes. In the absence of clustering, we observed a substantial error rate consistent with our results obtained for the single-plasmid controls.

**Table 5:** 10-plasmid controls preparation.

| Barcodes | 10P-A |  | 10P-D |  | 10P-G |  |
| --- | --- | --- | --- | --- | --- | --- |
|  | Plasmid copies | PCR cycles | Plasmid copies | PCR cycles | Plasmid copies | PCR cycles |
| TCACAGGGGTAA | 1.00E+05 | 29 | 1.00E+04 | 29 | 1.00E+05 | 22 |
| ACAAGACCGGAA | 1.00E+03 |  | 1.00E+04 |  | 1.00E+05 |  |
| ATATAGAGCTGT | 1.00E+03 |  | 1.00E+04 |  | 1.00E+04 |  |
| ACATACCTGCTA | 1.00E+02 |  | 1.00E+04 |  | 1.00E+04 |  |
| GTGTCAGGCACA | 1.00E+02 |  | 1.00E+04 |  | 1.00E+03 |  |
| TGCCACTCTAGC | 1.00E+01 |  | 1.00E+04 |  | 1.00E+03 |  |
| CTCGATTCACTC | 1.00E+01 |  | 1.00E+04 |  | 1.00E+02 |  |
| GAACCCGTGGAA | 1.00E+01 |  | 1.00E+04 |  | 1.00E+02 |  |
| CTGTATATTTTA | 1.00E+01 |  | 1.00E+04 |  | 1.00E+01 |  |
| GAAACCATGACA | 1.00E+01 |  | 1.00E+04 |  | 1.00E+01 |  |

The 10-plasmid controls allow us the resolution to experimentally compare different Starcode algorithms with a range of parameters for how well they control error. Starcode offers several clustering algorithms, including its default message-passing algorithm as well as a greedy “spherical” algorithm (14). Both algorithms use the count information, as well as a Levenshtein distance threshold. But rather than starting with the highest-count sequence as a greedy algorithm would, the message-passing algorithm begins with the lowest-count sequences and gradually builds up clusters taking relative count numbers into account. The centroid is defined as the representative barcode. Two clusters are combined into a larger cluster only if the centroids are within the threshold distance, and the total counts of one cluster are 5 times larger than the counts of the other. The condition on relative count sizes addresses the likelihood that an erroneous barcode will have far fewer reads than a true barcode. If two true barcodes happen to be within the distance threshold, more likely than not, their counts do not differ by more than a factor of 5. When applied to the 10-plasmid controls, we observed that the message-passing algorithm reduces noise at lower values of L as compared to the greedy spherical algorithm (supplementary Figure S4).

Applying different Levenshtein distances using Starcode’s message-passing algorithm to the 10-plasmid control series, we observe that distances of L=2 or L=3 substantially reduced errors in recall (Figures 4 and supplementary Figure S5). This raises the question of what Levenshtein distance to apply. The main downside of higher thresholds for L would be overclustering (or grouping separate true barcodes as the same). So what impact is L=3 likely to have on overclustering? We evaluated this question via computational simulation. Distances between 5000 random 12-mers were compared across each of 20 trials. Non-identical 12-mers within L=3 of each other never exceeded 0.074% of the total pairwise comparisons – less than a tenth of a percent. Thus, with the assumption that barcodes are random and the sequence space covered in the libraries is small, L=3 should have little effect on overclustering for our library.

**Figure 3:**
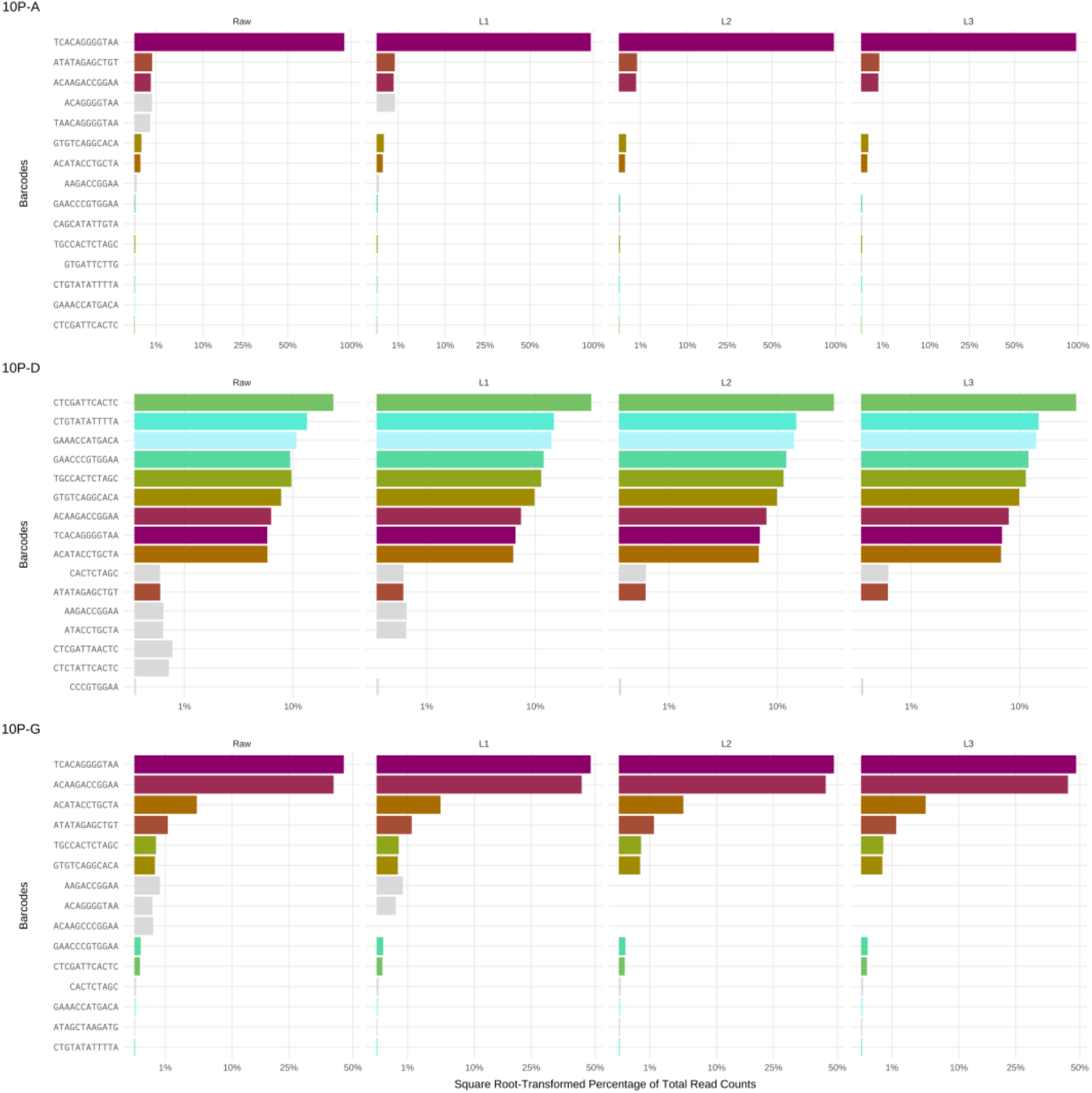
Illumina sequencing reads from 10-plasmid controls using different clustering distances. The y-axis depicts the barcode sequence; the x-axis shows the square root-transformed percentage of total read counts. The colored bars represent the control barcodes. Gray bars represent the most common erroneous barcodes within a library, across the Levenshtein distances. Here we show the 10-plasmid controls 10P-A, 10P-D and 10P-G. Additional 10-plasmid controls are shown in supplementary Figure S5.

**Figure 4:**
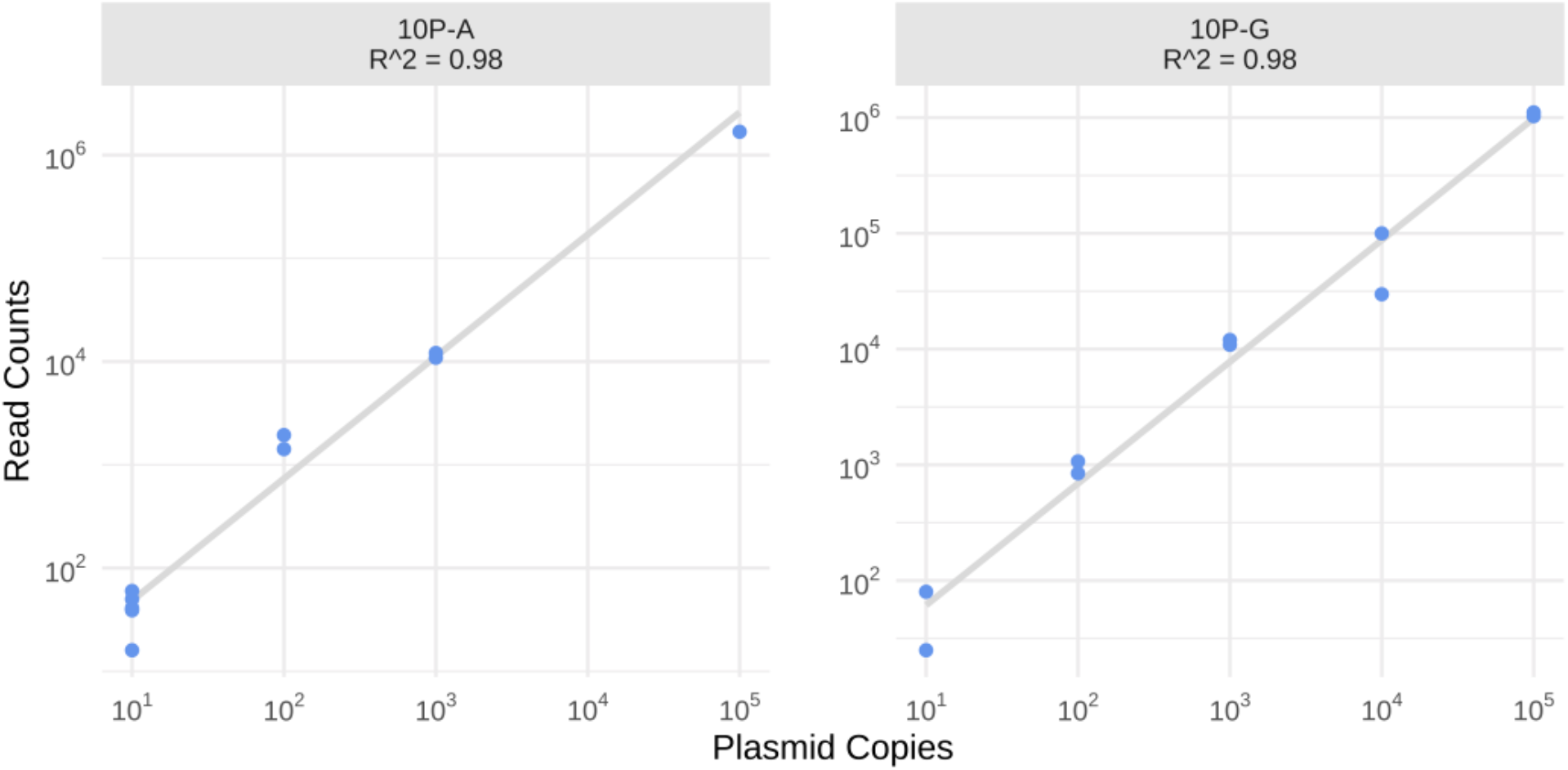
Linearity plots of 10-plasmid controls with L3 clustering parameter. The log10 transformed x-axis shows the copy number of plasmid inputs, the log10 transformed y-axis represents L3 clustered read counts. Linear regression trendlines are plotted in gray, with corresponding R^2^ values. Linearity of the 10-plasmid control series 10P-A and 10P-G is shown (the linearity of additional 10-plasmid controls is shown in supplementary Figure S6).

Finally, we also used the 10-plasmid control to probe for linearity of recall barcodes and to determine the limit of detection. Raw counts of the 10-plasmid controls show a linear recall of barcodes of at least three orders of magnitude with a sensitivity reaching as low as 10 input copies per enrichment PCR in most controls (Figure 4 and supplementary Figure S6).

From the analyses of these controls, we conclude: (i) Applying the message-passaging algorithm and a clustering distance of L=3 consolidated most of the variation with the true barcodes with little downside. (ii) Differences in PCR cycle amplification did little to affect the relative abundance of barcoded virus genomes. (iii) At least on a small number of variant barcodes with well-defined inputs, our approach provides a linear recall spanning a range of at least three orders of magnitude.

### Barcodes in the plasmid library

One of the steps for the preparation of a barcoded muPyV stock requires the generation of a pool of plasmids containing the barcoded virus genome (similar to plasmids in our known barcode controls above) that will be subsequently isolated from the vector, circularized and transfected into cells. In our experimental conditions, we estimated the pool of barcoded plasmids to contain approximately 5700 different true barcodes based on the estimated bacterial colonies number that make up the library. This is an unreasonable number to sequence by the Sanger method. However, understanding the barcode diversity in the plasmid library is an important prerequisite to understanding the diversity of our virus library. We therefore applied Illumina sequencing to the plasmid library.

### Clustering of the plasmid library

We compared the distribution of barcode ranks in the plasmid library (based on read abundance) between the raw reads and clustered reads using L=1, 2, and 3 (Figure 5). The curve of raw (unclustered) barcodes ordered by rank exhibits a shallow shoulder centered near 6000 barcodes but extending to 200,000 (note, the cutoff for the graph in Figure 5 is at 10,000 barcodes). This shoulder indicates a marked drop in abundance of reads per barcode, consistent with the less abundant barcodes resulting from error (note, the log scale used greatly inflates the apparent abundance of less common barcodes). If one decided to discard all presumed erroneous low abundance barcodes, it would be difficult to know exactly where to draw the cutoff threshold. As clustering progresses from L=1 to 2 to 3, a pronounced shoulder materializes just under 5000 barcodes, a number broadly compatible with the estimated size of the plasmid library.

**Figure 5:**
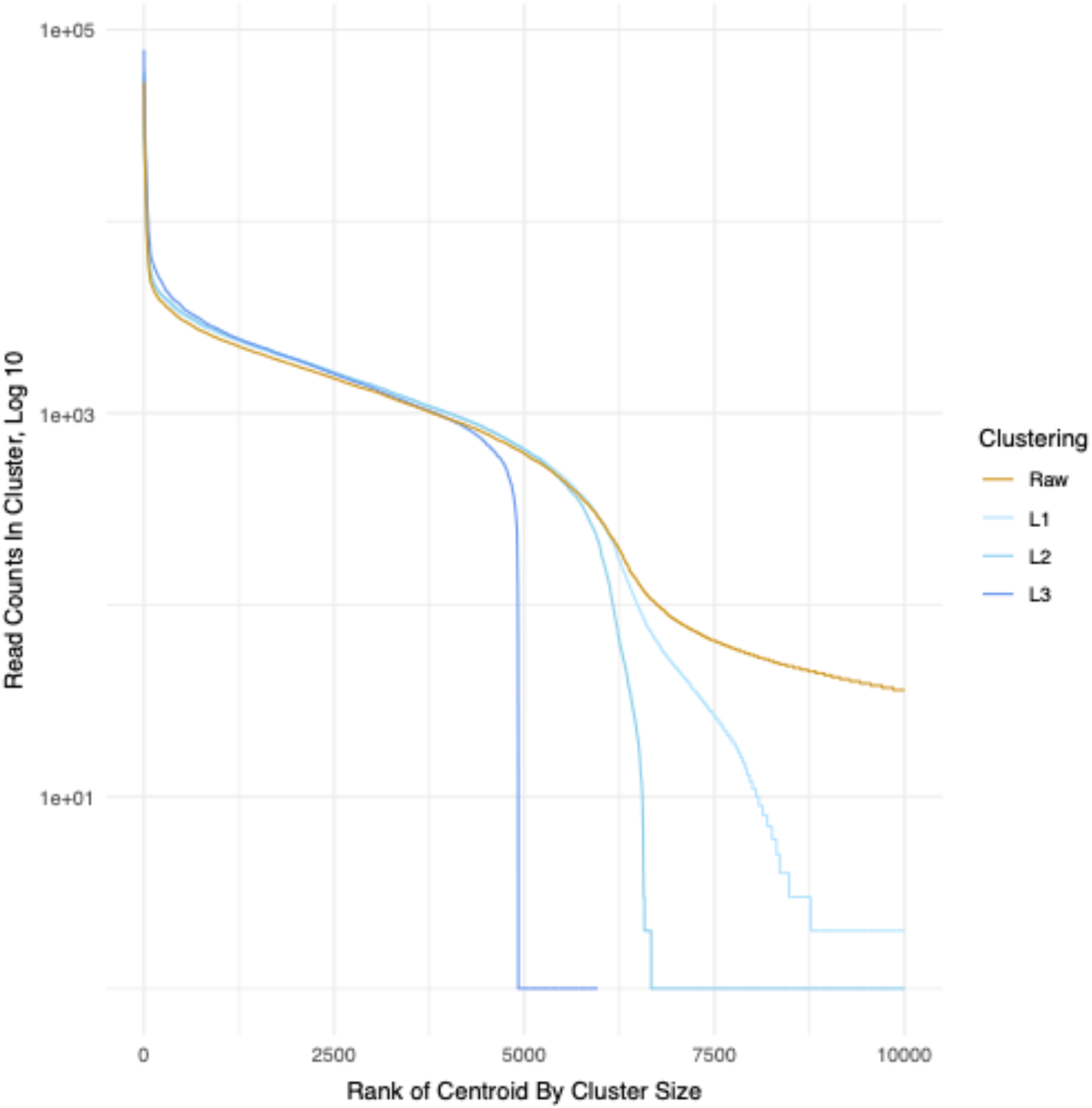
Comparison of barcode counts between raw and different clustering distances. The highest-count cluster is ranked 1. This figure shows that the number of barcodes that are called decreases with the increasing clustering distance; clustering with L=3 substantially decreases the barcode counts called in the raw sequencing reads. The figure is cut off to include only the 10,000 most abundant barcodes to focus on the “elbow” where the number of barcodes that are called display a steep drop-off.

After clustering with L=3, the fewest reads of any barcode numbered several hundred, and ∼ 3800 barcodes had at least 1000 reads (Figure 5). However, there is considerable variation in the number of reads across clustered barcodes, with approximately 100-fold more reads for rank 1 than for rank ∼ 4000. The distribution of read numbers for clustered barcodes broadly matches that for unclustered barcodes up to rank of 4000 (Figure 5), demonstrating this large variation is not an artifact of clustering.

Our results show that true barcodes in the plasmid pool are not evenly distributed across the pool with a ∼100-fold difference in abundance between the most common and least common cluster. For our series of plasmid controls, where we deliberately created differences in DNA abundance, these differences were faithfully reproduced in the Illumina sequencing reads (Figure 4). This suggests that these apparent large differences in our plasmid library are not introduced by sequencing artifacts. We suggest that this apparent large variability in plasmid counts arises due to differences in amplification and/or propagation of the plasmids.

Every cluster consists of several barcodes, one of which serves as the representative for the cluster (also known in as the “centroid” in the literature). To get a sense of how much the clustering contributes to the total abundance of the barcode compared to the centroid, Figure 6 shows the proportional contribution of the centroid to the cluster. We see that aside from some rare exceptions, centroids contribute the overwhelming majority of counts to their clusters. This is consistent with the expectation that clusters represent error-correction of what are likely the “true” barcodes, which are assumed to be the centroids.

**Figure 6:**
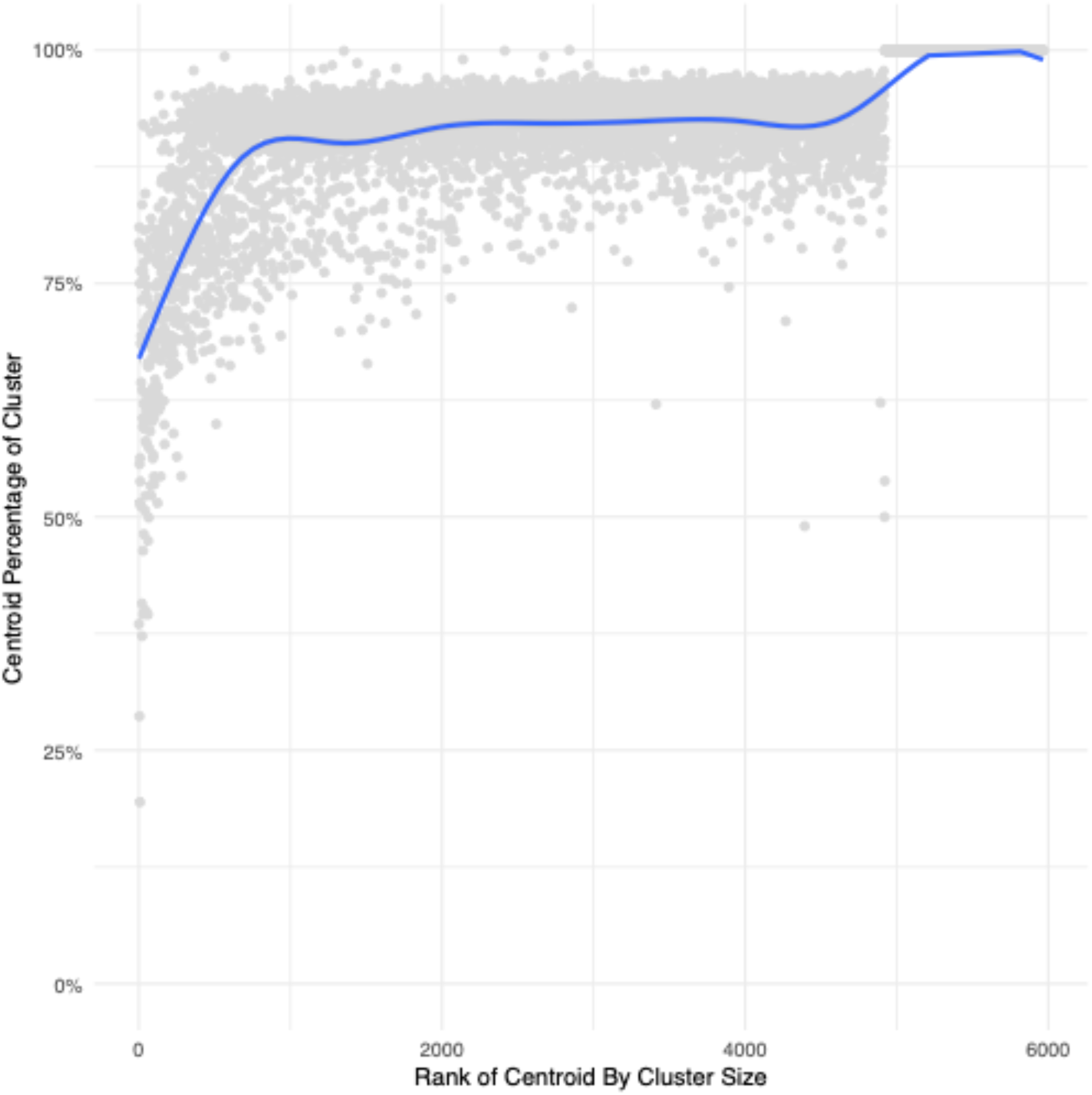
Centroid counts as percentage of cluster counts. Each gray dot represents the percentage of counts in a cluster (using L3 clustering) from the plasmid library that comes from its centroid. The centroids are ordered by the size of their cluster. The highest-count cluster is ranked 1. A smoothed curve (in blue) is fitted to the dot plot. This plot shows that the majority of cluster counts come from the defined centroid.

### Assigning a cutoff

As expected from the design of the library, the majority of barcode lengths recalled were 12 nt long (Figure 7). However, we also noted a small subfraction of barcode reads that had a median length of 36 nt, even after clustering. These were clearly not intended in our experimental design for construction of the library and were of very low abundance. We therefore applied a series of cumulative read count cutoffs (top 90%, top 99%, or top 99.9% most abundant (as described in Figure 1)) to weed out these low abundance likely artifacts. (A cutoff of 90% means that the remaining barcodes account for 90% of the total counts.) This analysis shows that all three cutoffs removed most of the aberrant size barcodes. The difference in the number of barcodes left after applying a 99% or 99.9% cutoff was small, and we therefore applied a conservative cutoff of the top 99% most abundant reads to subsequent work on the virus library. After applying the cutoff, barcodes of length 12 nt account for 90.6% of the total count. Barcodes of length 11 or 12 nt account for 95% of the total count.

**Figure 7:**
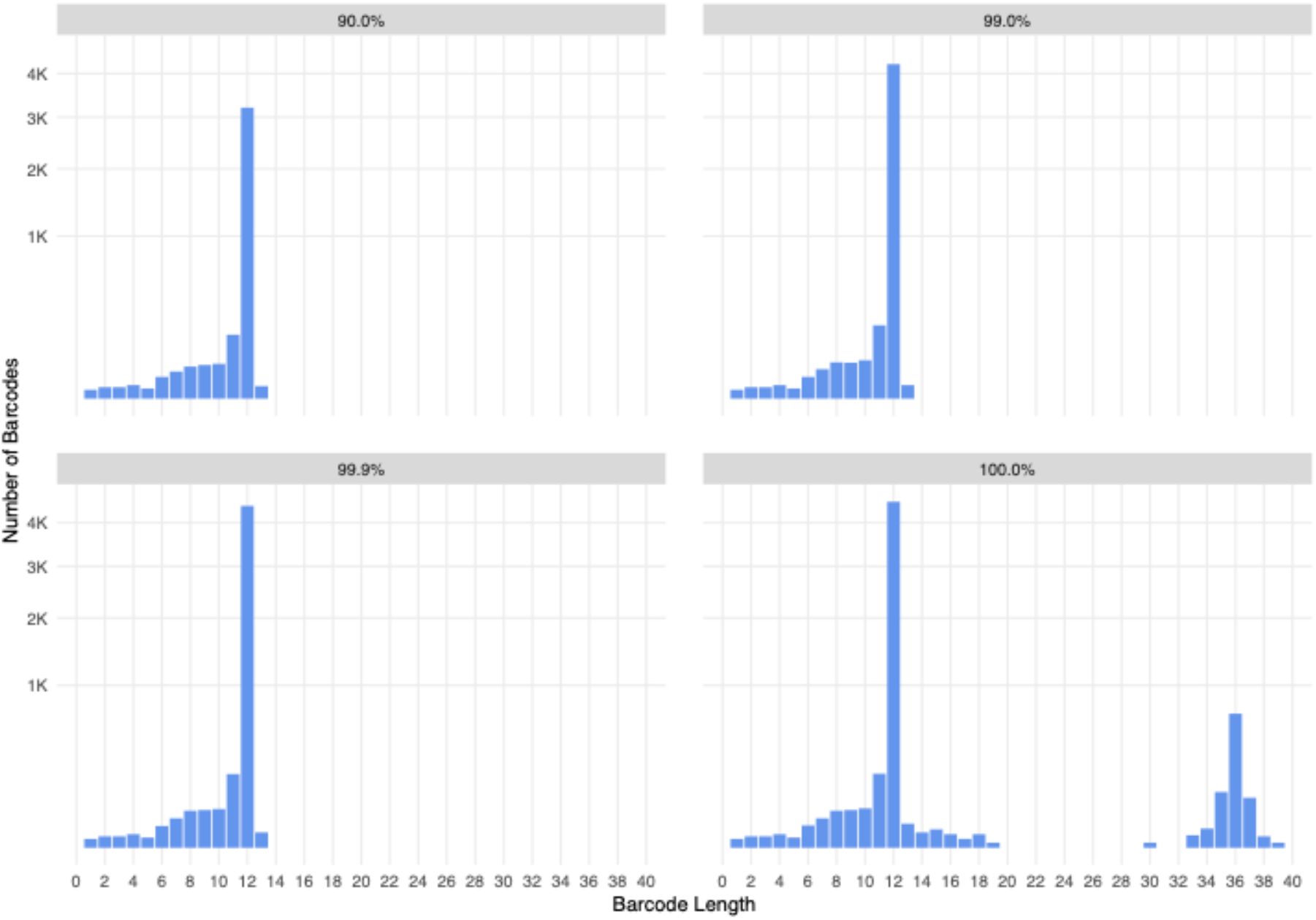
Barcode length distributions for 90%, 99%, and 99.9% sequencing reads cutoffs. Applying a cutoff for barcode abundance eliminates most recall of longer barcodes that were not intended in the original design for a 12-nucleotide barcode library. These barcodes are derived from the L3 clustering. The y-axis is square root-transformed so low values are more visible.

We evaluated barcode properties of our plasmid library after clustering and applying the 99% cutoff (Figure 8). The histogram of pairwise Levenshtein distances between barcodes shows a narrow peak at 8. This indicates that the barcodes are well-separated, which will help to properly assign barcodes in downstream experiments that may have a small number of errors associated with a true, parent barcode. The distribution of pairwise distances and size distribution are consistent with a randomly-generated barcode library that arose as expected from the experimental design.

**Figure 8:**
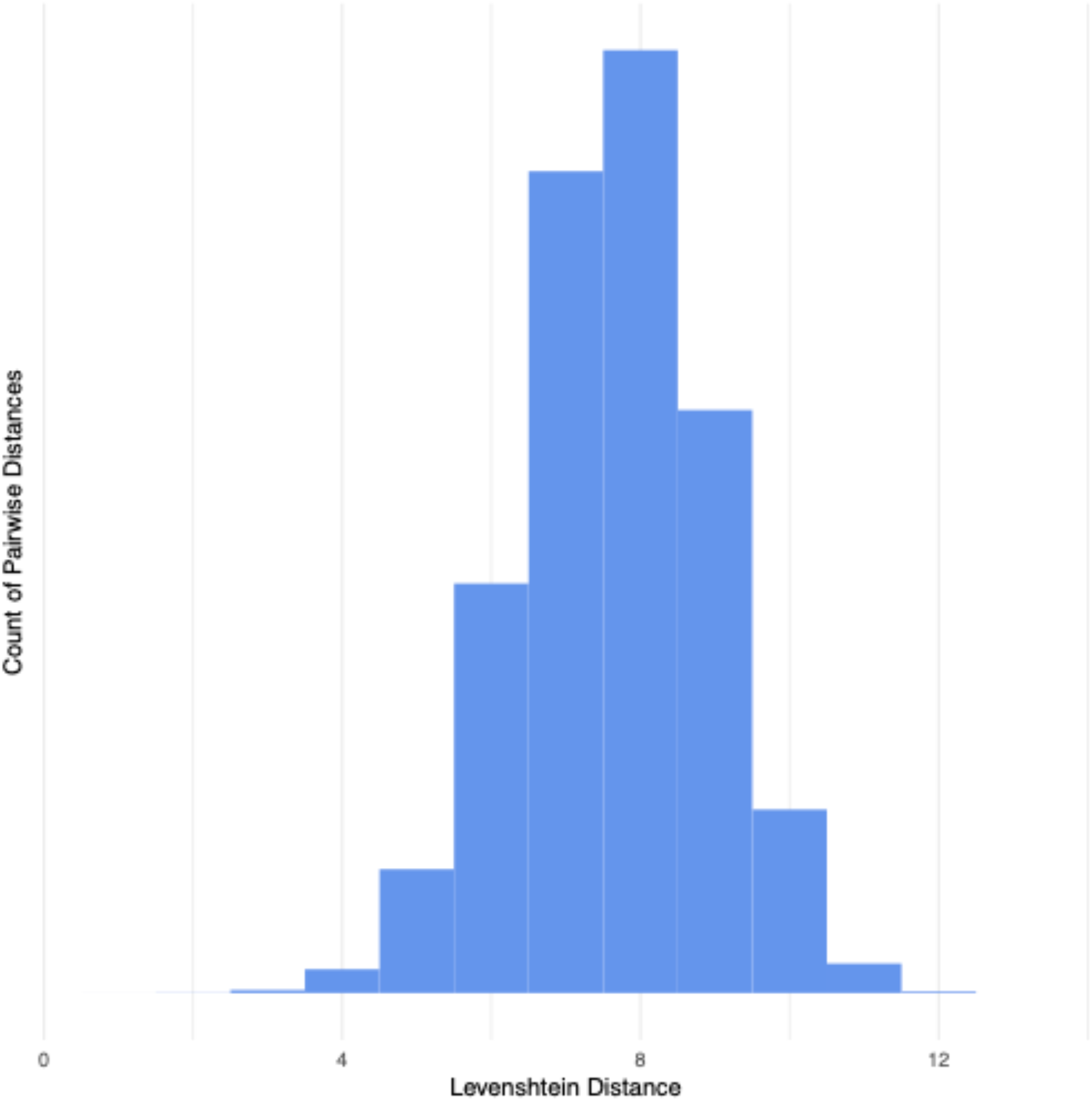
Distributions of the barcode pairwise distances within the plasmid library. The figure represents the pairwise Levenshtein distances between centroids in the plasmid library after applying a L3 clustering distance and a 99% reads cutoff. Data show that the proportion of possible barcode sequence space covered is sparse.

In summary, our analysis of the plasmid library shows (i) there is more variation in barcode abundance than intended in the original library design, (ii) that clustering works in eliminating the tail of low abundance (most likely error) reads, (iii) applying a 99%-most-abundant-reads cutoff eliminates most unintended aberrant sized inserts, and (iv) centroids are usually the dominant member of a cluster consistent with the clustering approach working as intended in identifying true barcode sequences.

### Clustering of barcodes in the virus library

We created the barcoded virus stock by transfecting cells with virus genomes that had been isolated from the plasmid library and circularized. To estimate the barcode composition of this library, we sequenced this library four times in two separate sequencing runs (one technical replicate of the virus library in the first sequencing run and three technical replicates in a second run) and for each we applied the message-passing algorithm with clustering Levenshtein distance of 3 with a 99% cutoff. We compared the overlap between the technical replicates and determined that most barcodes were recalled in all four replicates (Figure 9). The sizes of the 4 technical replicate libraries after clustering and cutoff were remarkably similar, varying from 3985 to 4001 barcodes. Comparison of the four clustered virus libraries show 3555 barcodes shared across all 4 libraries, with another 331 shared across 3 libraries. This overlap is stark, with nearly all counts occurring in intersection of all 4 replicates. Using this method provides an estimate of approximately 3600 high confidence barcodes.

**Figure 9:**
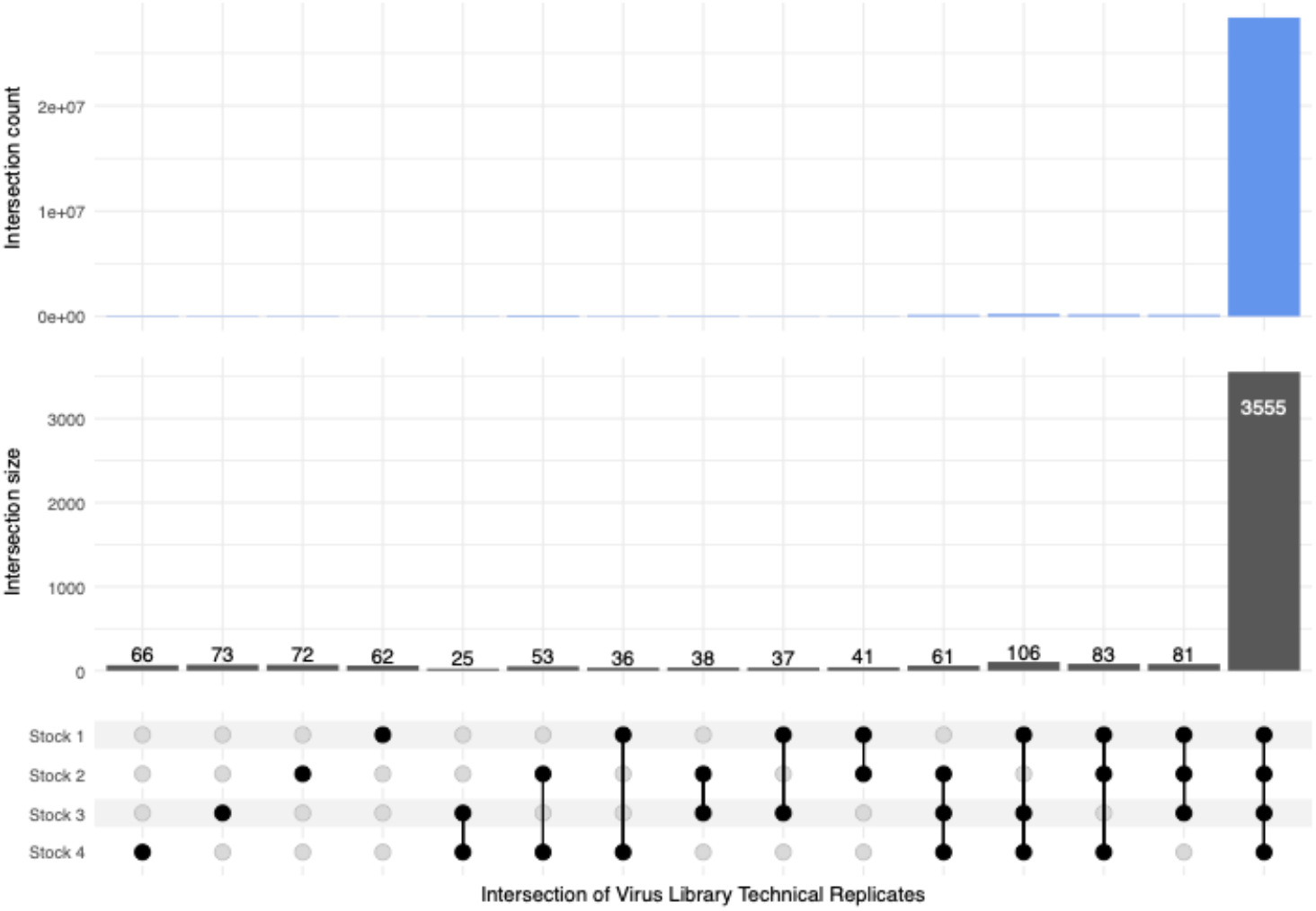
Technical replicates of the sequencing of the virus library. One replicate of the virus library was sequenced via Illumina sequencing and three additional technical replicates were sequenced in a separate run. The UpSet plot shows the number of barcodes that intersect amongst the four virus library technical replicates (lower panel), as well as the total counts of those barcodes in blue (upper panel). The libraries were clustered using L3 distance and a 99% reads cutoff was applied. The four replicates show a large degree of overlap in clustered barcodes.

To determine potential bottlenecks in the process of generating the barcoded virus library, we compared the overlap between the virus, plasmid, and control ligation libraries that were used to give rise to the virus library. After clustering and cutoff, the majority (90%) of the virus barcodes from the pool of our 4 replicates of the virus library were recalled in both our sequencing of the plasmid libraries and the control ligated virus genomes, with 5% of the virus barcodes being unique to the virus library. Examining the counts, 95% of the counts for the pool of the virus library are for barcodes that are also found in both the plasmid library and ligated virus genomes, and only 3% of the counts were unique to the virus library barcodes (Figure 10). We note that these numbers compare centroids of clusters; the similarity of the libraries may in fact be even higher, as the barcode chosen as the centroid of a cluster may differ between libraries, even if the clusters themselves are very similar. We conclude that the vast majority of barcodes present in the plasmid library are incorporated into the virus library. Thus, as would be predicted, a greater repertoire of barcodes could be obtained in the virus library if plasmid libraries are derived from an increased number of unique bacterial colonies.

**Figure 10:**
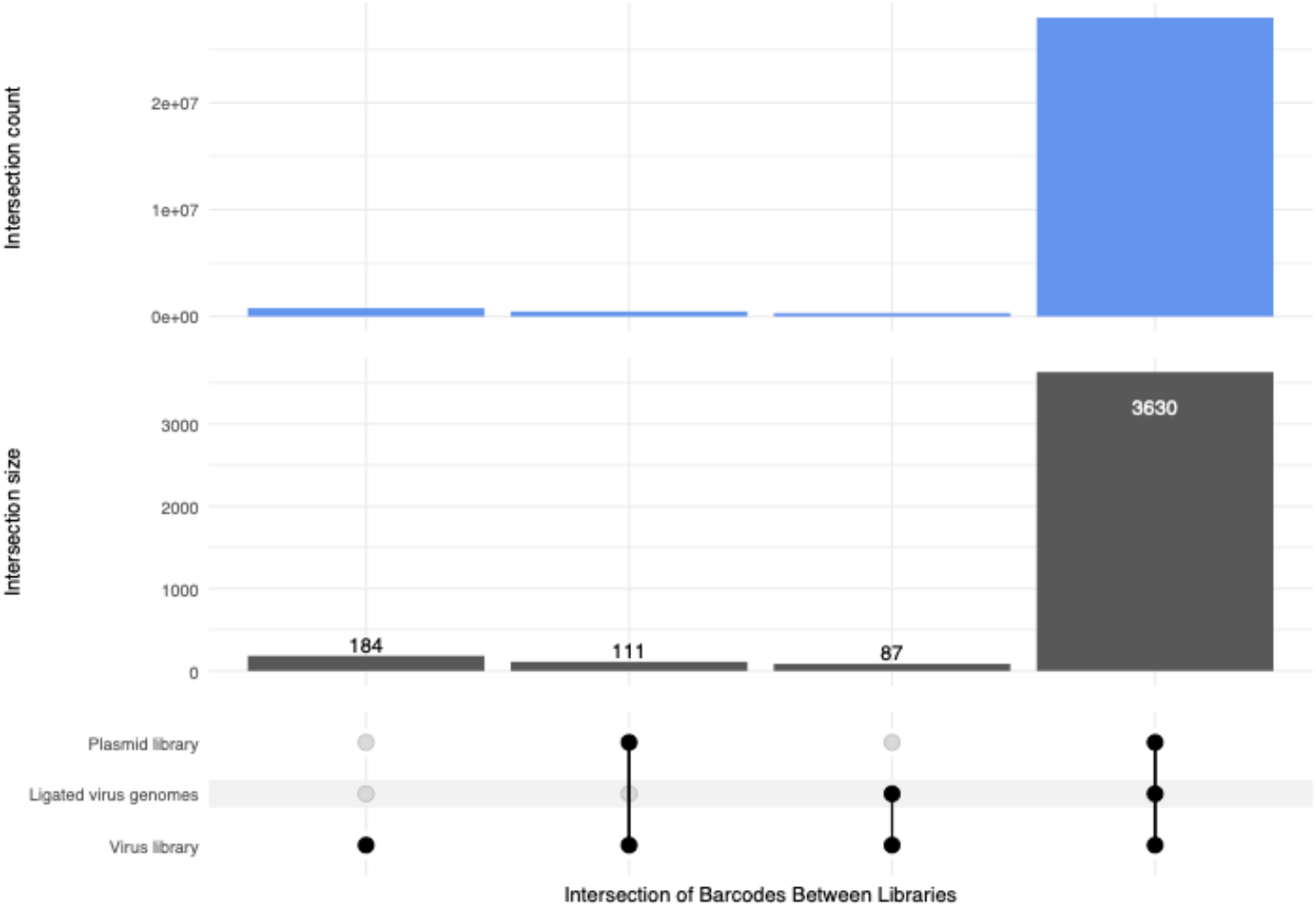
Overlap of clustered barcodes from the plasmid library, ligated virus genomes, and the virus library. L3 clustering distance along with a 99% reads cutoff was applied to the three libraries. In the lower panel, the UpSet plot shows the number of barcodes from the virus library that intersect with barcodes from the plasmid library and/or the ligated virus genomes. Data show that the vast majority of barcodes from the virus library are present in the plasmid library and the ligated virus genomes. In the top panel, the UpSet plot represents the total read counts associated with barcodes in the virus library that intersect with the plasmid library and the ligated virus genomes. Data show that the barcodes in all three libraries account for the overwhelming number of counts in the virus library.

After clustering and applying a 99% cutoff most barcodes were the expected size, we noted that approximately 9% of barcodes were less than 12 nts (ranging from 0-11 nucleotides in length). We determined that 88% of these shorter barcodes were also present in the plasmid library and ligated virus genomes that gave rise to the virus stock (Figure 11). Combined, these findings do not support overt pressure for viruses to lose the barcode under our experimental conditions and suggest that while some shorter barcodes may arise during virus propagation, shorter-than-expected barcodes mainly arise due to synthesis errors in the primers used to generate the barcoded plasmid libraries and/or deletions that occurred during production/propagation of the plasmids.

**Figure 11:**
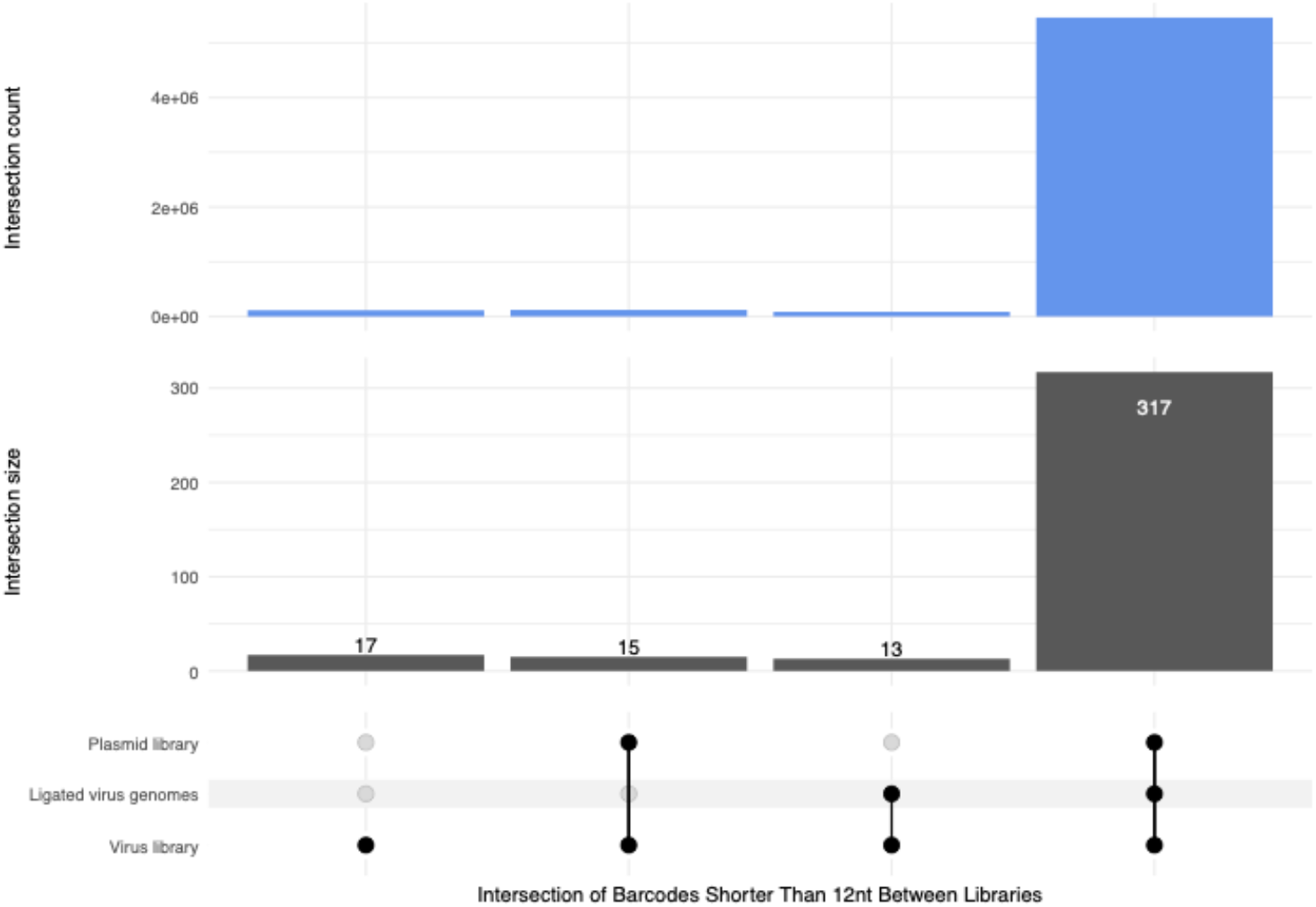
Overlap of clustered barcodes that are shorter than 12 nucleotides from the plasmid library, ligated virus genomes, and the virus library. L3 clustering distance along with a 99% reads cutoff was applied to the three libraries. In the lower panel, the UpSet plot shows the number of barcodes from the virus library that intersect with barcodes from the plasmid library and/or the ligated virus genomes. Data show that the majority of short (<12nt) barcodes in the virus library are also present in the plasmid library and ligated virus genomes. In the top panel, the UpSet plot represents the total read counts associated with shorter barcodes in the virus library that intersect with the plasmid library and the ligated virus genomes. Data show that the shorter barcodes that overlap in all three libraries account for the majority of shorter barcodes in the virus library, suggesting that they did not arise during the course of infection.

Given that the plasmid library had large differences in the relative abundance of its members, we asked if the virus library barcode abundance was reflective of this. We observe differences in individual barcode abundance in the virus library ranging between 1,544 counts for the least abundant up to 243,503 counts for the most abundant barcode. There is good correlation between the abundance of plasmid and virus library barcodes, with a Pearson correlation of 0.87 (Figure 12). Combining the above findings, these results suggest that, after the original transformation of bacteria, there were no major bottlenecks in generating virus, with the virus library largely reflective of the composition of the plasmid library used to give rise to it.

**Figure 12:**
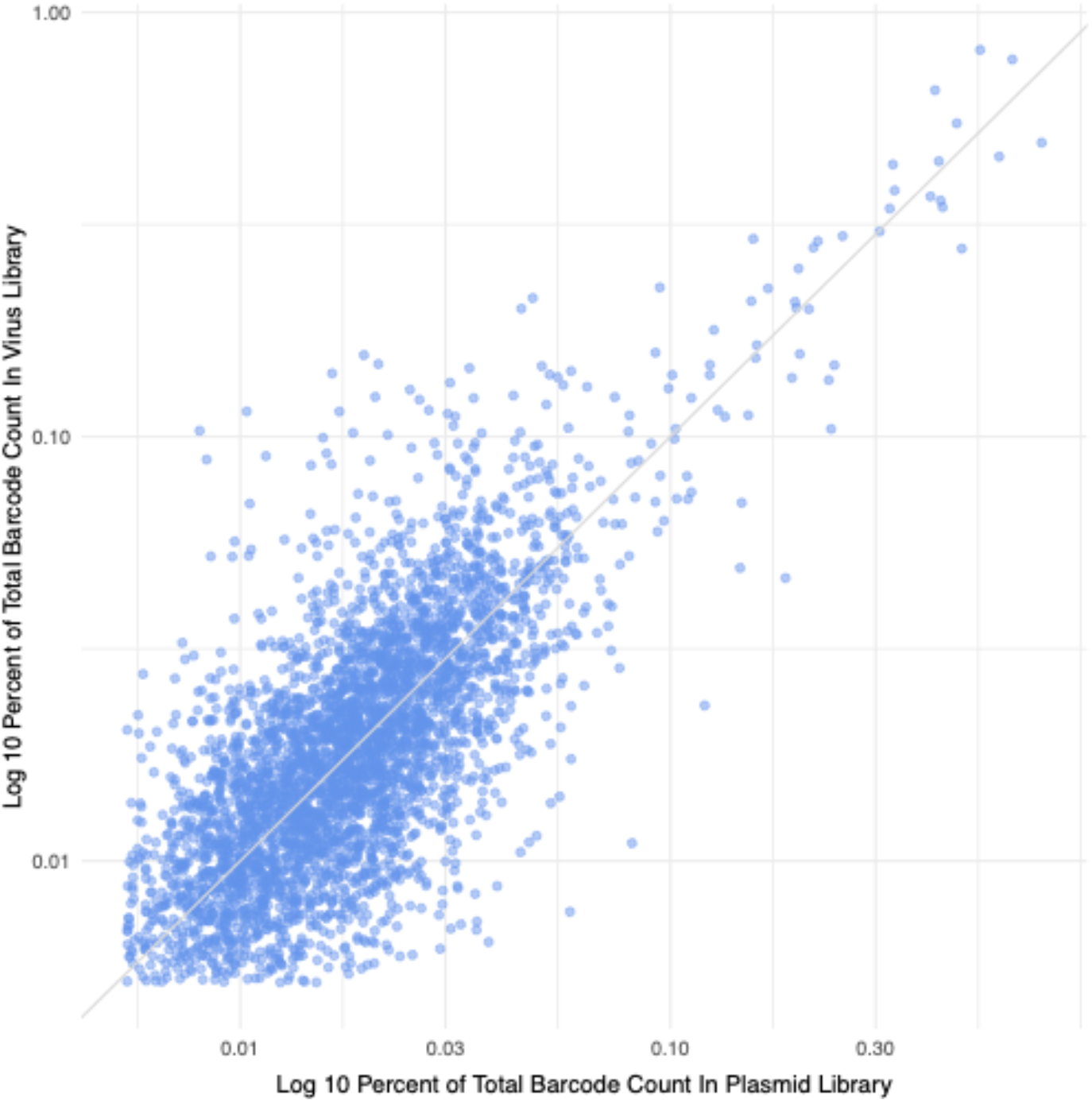
Abundance of clustered barcodes in the plasmid and virus libraries. For each barcode its log10 percentage of counts in the plasmid library is plotted on the x-axis, and its log10 percentage of counts in the virus library is plotted on the y-axis. A trendline with slope 1 through the origin is plotted. Data show a correlation with the more abundant barcodes in the plasmid library generally giving rise to the more abundant barcodes in the virus stock.

In summary, (i) clustering under our conditions of the virus stock results in highly similar clusters between technical replicates, supporting the good reproducibility of our approach, (ii) there is a large agreement of clusters between the virus and plasmid libraries both in identity and abundance, and (iii) under the conditions we applied, after generation of the plasmids that give rise to the virus library, the virus library was not subject to major bottlenecks during its generation. We conclude that the wet bench and computational approaches developed here are suitable for tracking longitudinal parallel infections with barcoded microbes.

## DISCUSSION

Conducting parallel infections with multiple variants of a pathogen is a longstanding way to assay competitive fitness advantages as well as to track the dynamics of infection(8,9,19–25). Studies that employ genetically tagged viruses can track the comparative progression of infection based on scoring differences in abundance of each variant. In its simplest form, this approach can be applied to as few as two differentially marked pathogens (24–28). The availability of large scale massively parallel sequencing greatly expands the number of potential differentially marked pathogens that can be assayed in a single experiment. Indeed, recent studies using RNA viruses or retroviruses have employed this technique to study a variety of aspects of infection (3,5,6,29–31). However, basic questions about how best to employ this approach, such as library generation strategy and the optimal computational methods to interpret sequencing data remain poorly characterized.

Here we describe the construction of a barcoded muPyV (murine polyomavirus) library. We estimate the library to be comprised of approximately 5000 different barcodes. Our data reveal a higher-than-expected error rate in our Illumina sequencing, which we speculate that despite our efforts using typical offset multiplexing primers and spiked PhiX DNA may be due the inherent low complexity of barcoded amplicons (18). The error rate of raw reads was such that it was difficult to determine with precision the exact number of true barcodes present in our virus library. This problem would likely be encountered in similar barcode approaches for other microbes. To solve this problem, we develop an approach based on extensive wet bench controls and computational simulation and analyses. We expect this approach to be informative to and/or reduce the workload of future studies that will utilize barcoded microbes.

The ratio of the most abundant to least abundant barcode in the virus library was greater than about 150-fold. This likely occurred because the barcode representation in our plasmid library that gave rise to the virus library spanned a similar range. This fortuitously allowed us to conclude that the ratio of virus barcodes tracks almost linearly to the barcodes in the input plasmids, which might be useful for those wishing to purposely generate libraries with different barcodes spanning a range of input copy numbers.

By examining different ratios of absolute amounts of plasmids containing known barcoded genomes, we were able to optimize clustering parameters to ensure accurate recall without loss of barcode resolution. By applying different PCR amplification cycles we also determined a range of PCR cycle conditions that did not result in substantial recall of incorrect barcodes. These conditions support accurate recall spanning at least three orders of magnitude of different amounts of barcode and sensitive to approximately 10 copies of barcode per reaction. Combined, using a prototypic model virus, we have developed a molecular and computational methodological approach we expect applicable to more broadly studying the dynamics of pathogen infection.

Consistent with others’ work showing that Illumina-based sequencing of amplicon libraries can have particularly high error rate even when Q value quality scores are high (>30) (18), we had a higher error rate (3.71 to 6.11%) as compared to other Illumina applications (typically ∼1%). We conclude that appropriate clustering is essential to interpreting the composition of a randomly generated barcoded virus library.

Many common clustering algorithms for sequences use top-down “greedy” algorithms, where the highest count or longest sequence is considered first, and all sequences within some threshold distance are made a cluster with the initial sequence as the representative, or “centroid”. The popular CD-HIT clustering software (32) works in such a way: the longest sequence is considered first, with all sequences within a threshold distance clustered with that sequence. Then the next-longest unclustered sequence is considered, and the process repeats. In applications where sequence lengths differ considerably, and longer sequences are generally considered higher-quality, this is a reasonable approach. In the case of barcodes, the situation is different. Our barcoding approach aims for consistent lengths, in this case 12-mers. Rather than large variation in length of sequence, we observe large variation in counts, spanning orders of magnitude. For this situation, which we expect would be encountered by others wishing to utilize nucleic acid barcoding, we demonstrate that using the Starcode message-passing clustering algorithm with Levenshtein distance of L=3 is appropriate.

We note that our approach to clustering is appropriate for determining the composition of the “true barcodes” of the initial inoculum (virus stock). However, performing the same clustering of reads from infected experimental samples is likely unhelpful. Indeed, the dynamics of a virus infection may dramatically impact the barcode counts, and the centroids of some clusters, especially low abundance true barcodes, may not be consistent between samples. Therefore, our results suggest an approach where first the “true” barcodes are identified in the initial virus library (as described in this work). Then, counts from infected experimental samples should be associated with these “true” barcodes. This could be accomplished by grouping with the nearest true barcode identified in the virus stock, where nearest is determined by Levenshtein distance, or more simply by counting only experimental sample barcodes that share sequence identity with true barcodes identified in the original virus stock.

Our combined analysis of the virus library, plasmid library that generated the virus stock, and various plasmid controls containing individual virus genomes with known barcodes, elucidated an effective strategy for generating and analyzing barcoded virus libraries.

The overall lessons learned from our efforts include:

i. Using long enough barcodes (in our case 12 base pairs), such that the number of actual barcodes is substantially less than the theoretical possible combinations, is desirable. In our case, the virus library comprised of approximately 5000 barcodes out of 1.68E+07 possible barcode sequence space. This allows for employing clustering without high risk of overclustering (the lumping of multiple, distinct true barcodes as a single barcode).
ii. Clustering of a reference virus stock is necessary and our empirical evidence suggests that using a message-passing algorithm may enhance performance for small length barcode (e.g., 12 nucleotide) microbe libraries. Although other virus barcode studies have employed clustering, we experimentally validated our parameters using Sanger sequenced plasmid controls to empirically estimate error rate, linearity, and functional clustering conditions. This afforded us confidence in the ability of our approach to determine relative barcode abundance.
iii. Even after clustering, some aberrant longer length barcodes with low counts remained in the library, which we eliminated by employing a read copy number cutoff, keeping the barcodes that accounted for 99% of the total counts.

In conclusion, we generated a library of a barcoded DNA virus. We describe unexpected outcomes of different steps of the construction of this library, devised control studies to explore optimal conditions for interpreting Illumina sequencing of barcoded pathogen libraries, and provide a validated wet bench and computational approach that may be useful to others wishing to employ barcoded microorganisms.

## AVAILABILITY

Code used to analyze data, for simulations, and to generate figures is available in the GitHub repository: https://github.com/bennigoetz/implementing-barcodes-dna-virus

## ACCESSION NUMBERS

Raw FASTQ files of Illumina sequencing data are deposited in the NCBI Short Read Archive (SRA) under the accession number PRJNA791340. They are available at: https://www.ncbi.nlm.nih.gov/sra/PRJNA791340

## SUPPLEMENTARY DATA

Supplementary Data are available at Plos Computational Biology online.

## AUTHOR CONTRIBUTIONS

Designed experiments: SB, JJB, CSS

Performed experiments: SB

Formatted data and figure compilation: SB, BMG

Generated Illumina libraries: SB

Developed and applied computational approaches: BMG

Interpreted data: SB, JJB, CSS, BMG

Managed project and procured funding: CSS

All authors assisted in the writing and editing of the manuscript.

## ACKNOWLEDGEMENT

The authors would like to thank Rodney P. Kincaid for helpful discussions regarding preparation of the Illumina library and strategies for processing the raw reads, Bob Garcea (University of Colorado at Boulder) for providing antibody against muPyV capsid protein and Aron Lukacher (The Pennsylvania State University Hershey) for providing NMuMG cells and helpful discussions.

## SUPPLEMENTAL MATERIALS

### SUPPLEMENTARY DATA

**Table S1.** 10-plasmid controls preparation.

| Barcodes | 10P-B |  | 10P-C |  | 10P-E |  | 10P-F |  |
| --- | --- | --- | --- | --- | --- | --- | --- | --- |
|  | Plasmid copies | PCR cycles | Plasmid copies | PCR cycles | Plasmid copies | PCR cycles | Plasmid copies | PCR cycles |
| TCACAGGGGTAA | 1.00E+04 | 32 | 1.00E+03 | 35 | 1.00E+04 | 29 | 1.00E+05 | 29 |
| ACAAGACCGGAA | 1.00E+04 |  | 1.00E+03 |  | 1.00E+04 |  | 1.00E+05 |  |
| ATATAGAGCTGT | 1.00E+02 |  | 1.00E+03 |  | 1.00E+04 |  | 1.00E+04 |  |
| ACATACCTGCTA | 1.00E+02 |  | 1.00E+01 |  | 1.00E+04 |  | 1.00E+04 |  |
| GTGTCAGGCACA | 1.00E+01 |  | 1.00E+01 |  | 1.00E+04 |  | 1.00E+03 |  |
| TGCCACTCTAGC | 1.00E+01 |  | 1.00E+01 |  | 1.00E+04 |  | 1.00E+03 |  |
| CTCGATTCACTC | 1.00E+01 |  | 1.00E+01 |  | 1.00E+04 |  | 1.00E+02 |  |
| GAACCCGTGGAA | 1.00E+01 |  | 1.00E+01 |  | 1.00E+04 |  | 1.00E+02 |  |
| CTGTATATTTTA | 1.00E+01 |  | 1.00E+01 |  | 1.00E+01 |  | 1.00E+01 |  |
| GAAACCATGACA | 1.00E+01 |  | 1.00E+01 |  | 1.00E+01 |  | 1.00E+01 |  |

**Figure S1:**
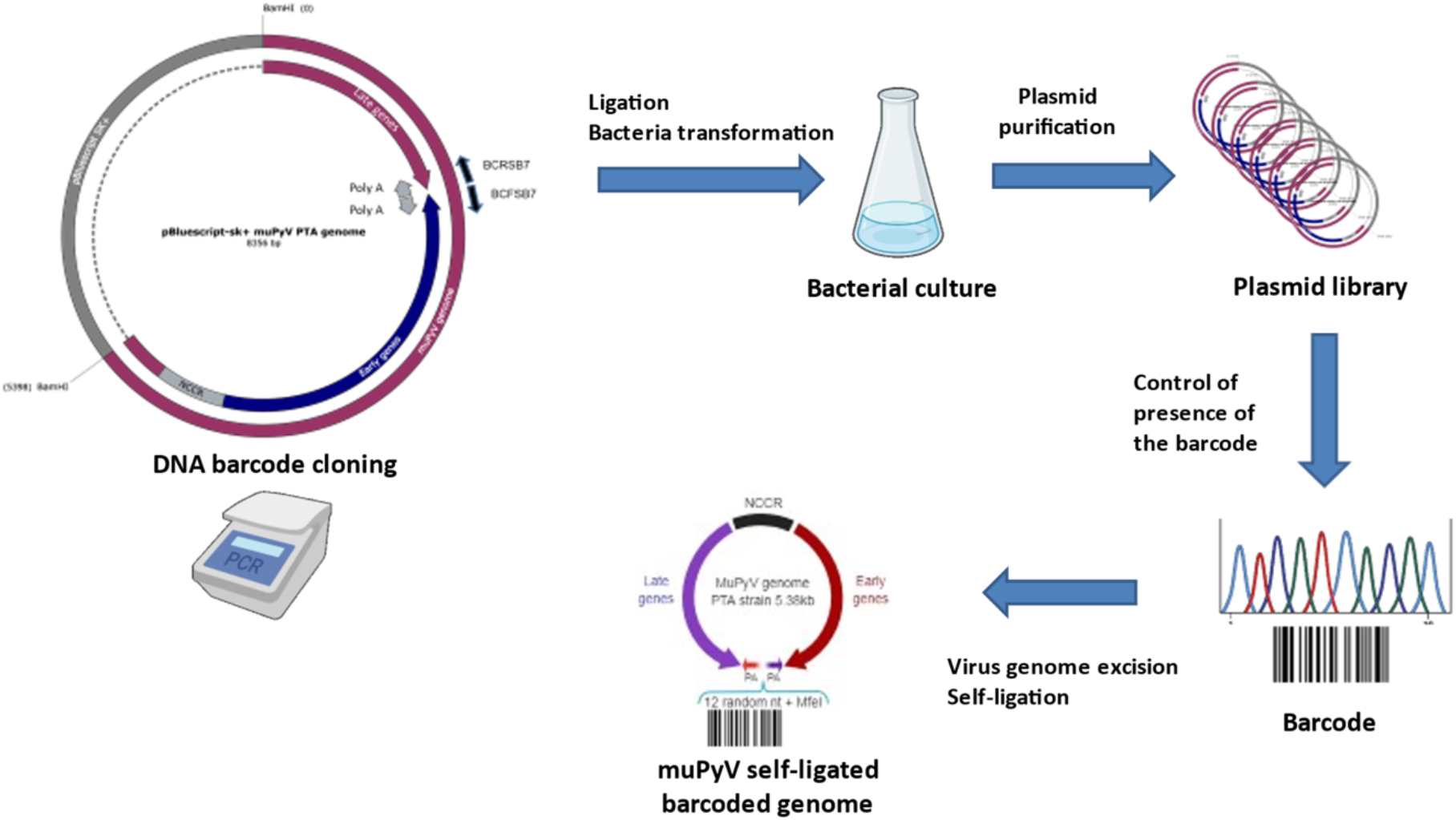
Schema representing the steps involved in the construction of the plasmid library and the barcoded muPyV genomes used for the subsequent generation of the virus library. Figure partly created with BioRender.com and SnapGene^®^ software (from Insightful Science; available at snapgene.com).

**Figure S2:**
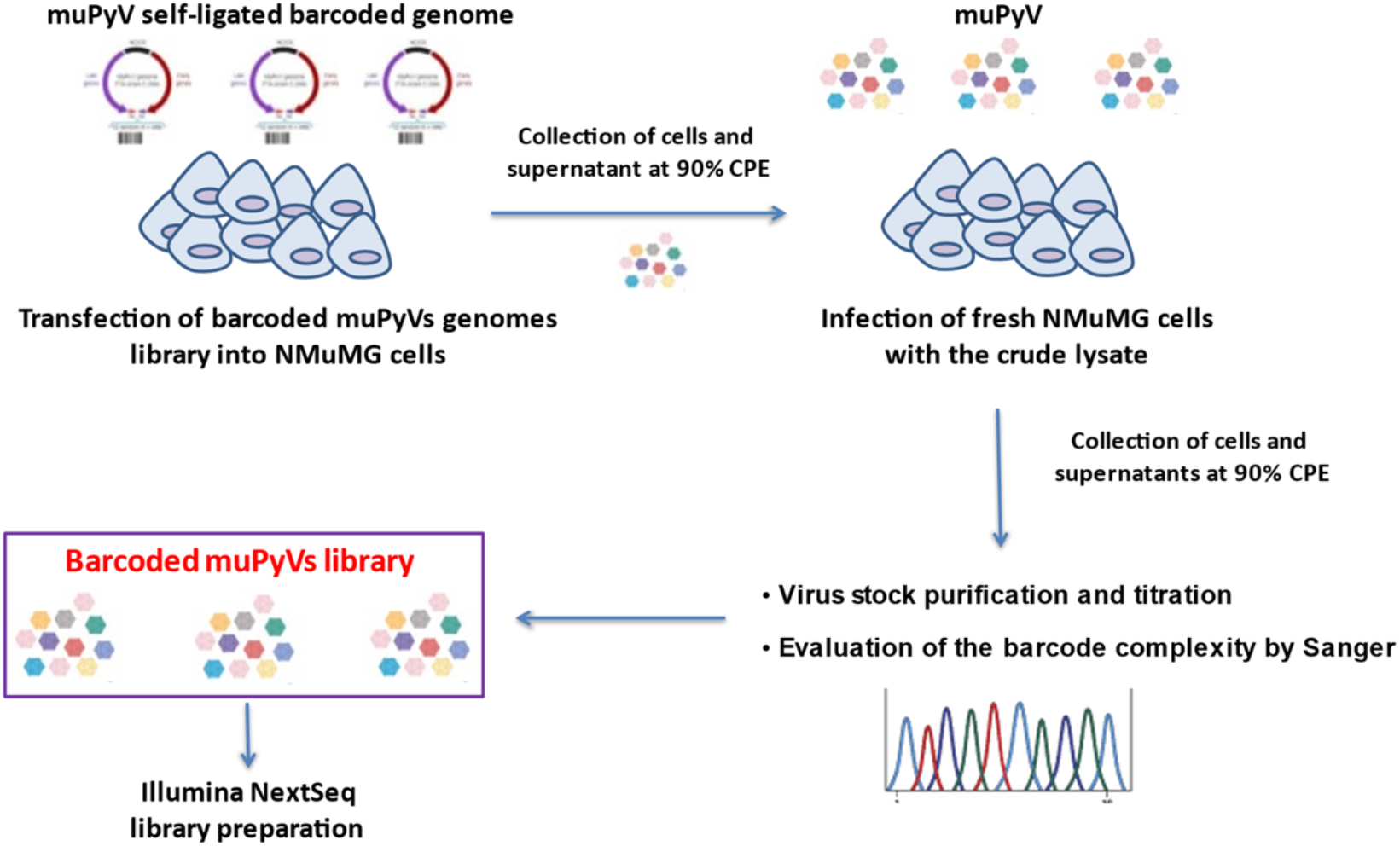
Schema representing the steps involved in the generation of the virus library. Figure partly created with BioRender.com.

**Figure S3:**
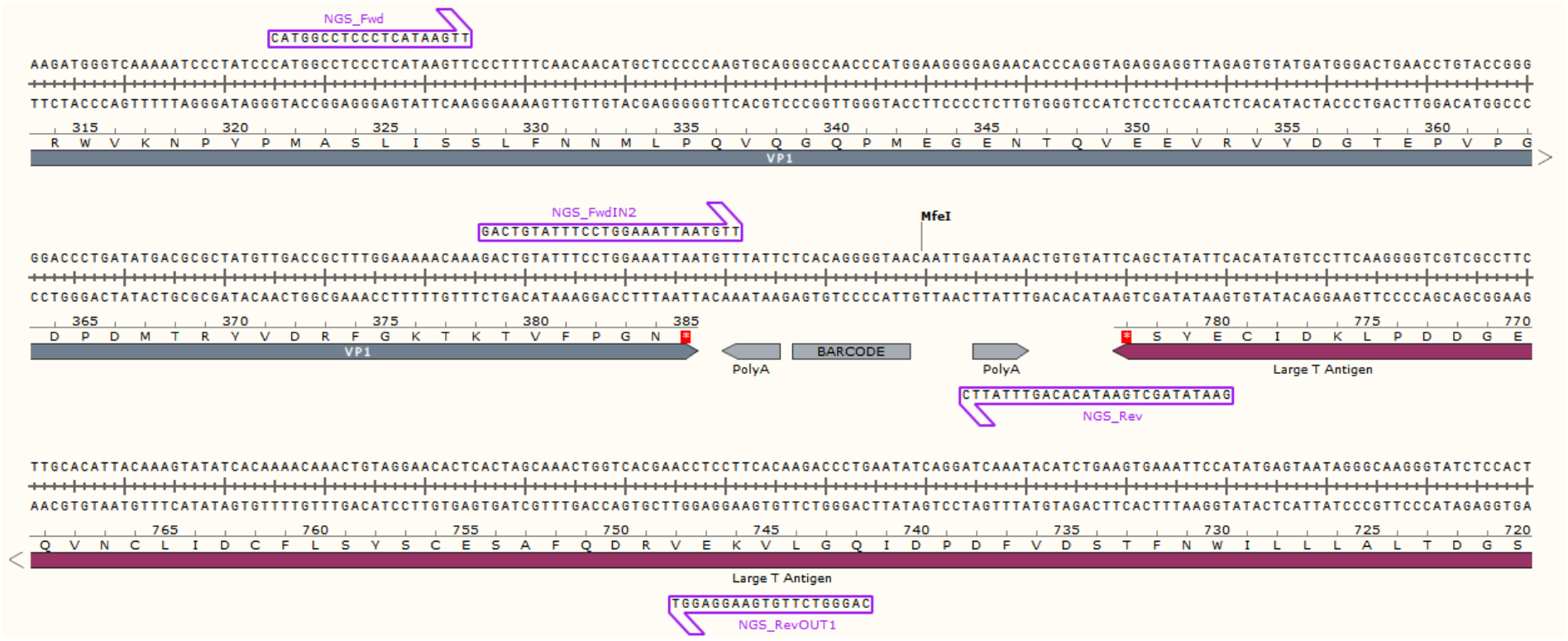
Schema representing the enrichment and indexing PCR primers binding site in the region surrounding the barcode in a muPyV barcoded genome. Figure partly created with SnapGene^®^ software (from Insightful Science; available at snapgene.com).

**Figure S4.**
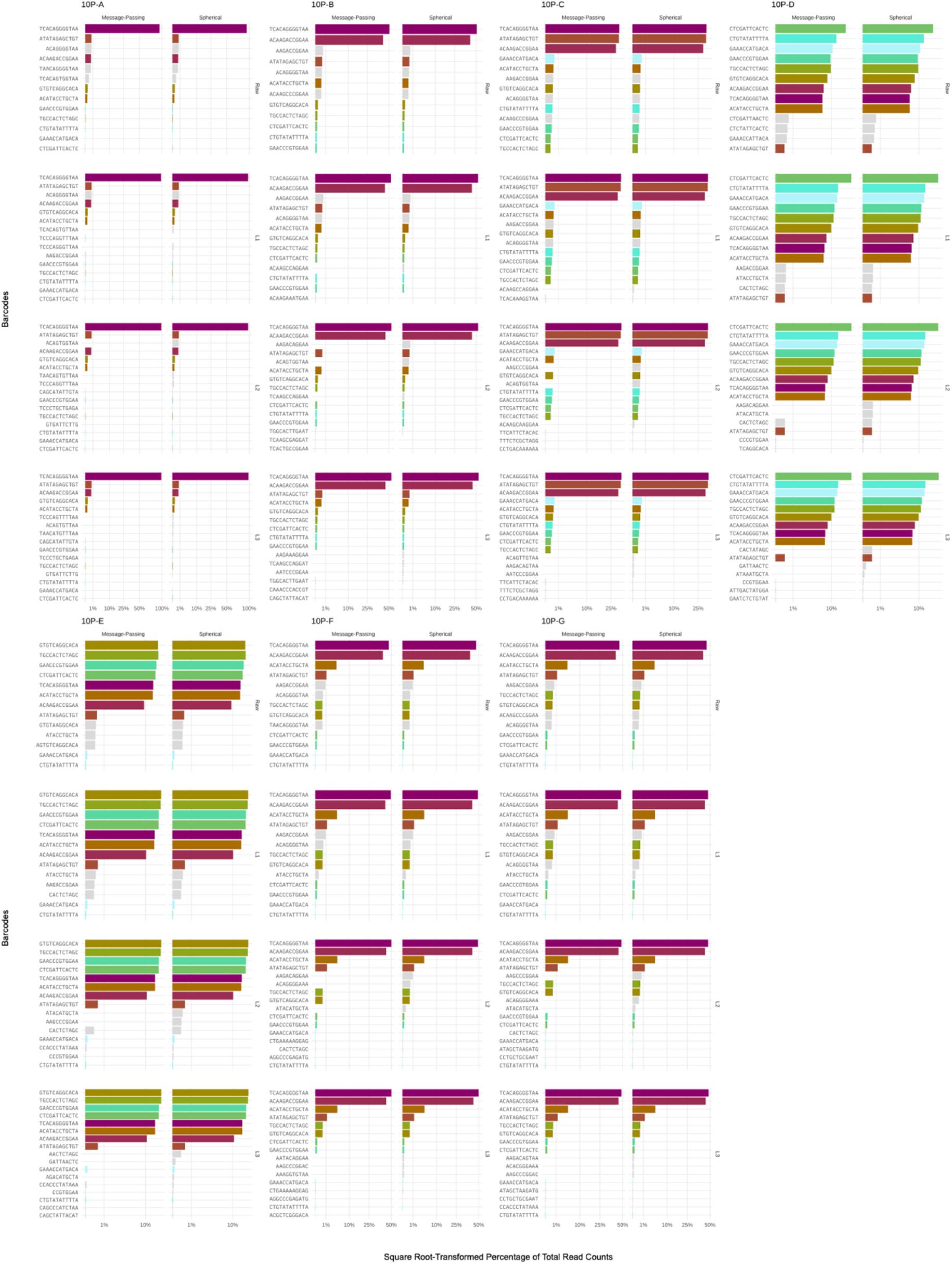
Illumina sequencing reads from 10-plasmid controls comparing Starcode’s message-passing vs spherical algorithms for increasing L distances. The y-axis panels vary the Levenshtein distance parameter; the x-axis panels show the message-passing vs spherical results side-by-side. The bar lengths are the square root-transformed total read count percentages. The colored bars represent correctly recalled input barcodes. Gray bars represent the most common erroneous barcodes for each L distance across either algorithm. Here we show the 10-plasmid controls 10P-A, 10P-B, 10P-C, 10P-E, and 10P-F.

**Figure S5:**
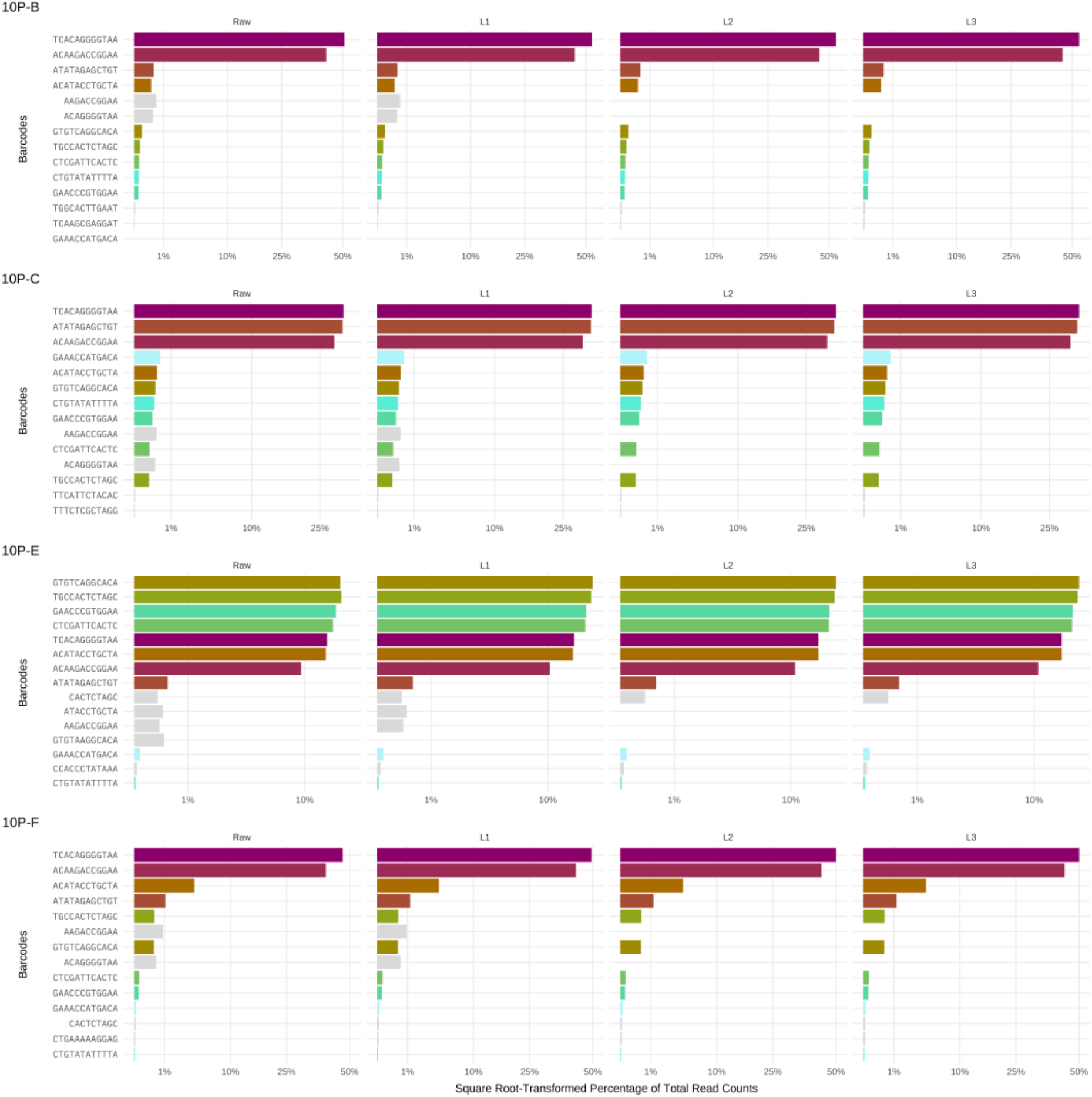
Illumina sequencing reads from 10-plasmid controls using different clustering distances. The y-axis depicts the barcode sequence; the x-axis shows the square root-transformed percentage of total read counts. The colored bars represent correctly recalled input barcodes. Gray bars represent the most common erroneous barcodes for each clustering parameter. Here we show the 10-plasmid controls 10P-B, 10P-C, 10P-E and 10P-F.

**Figure S6:**
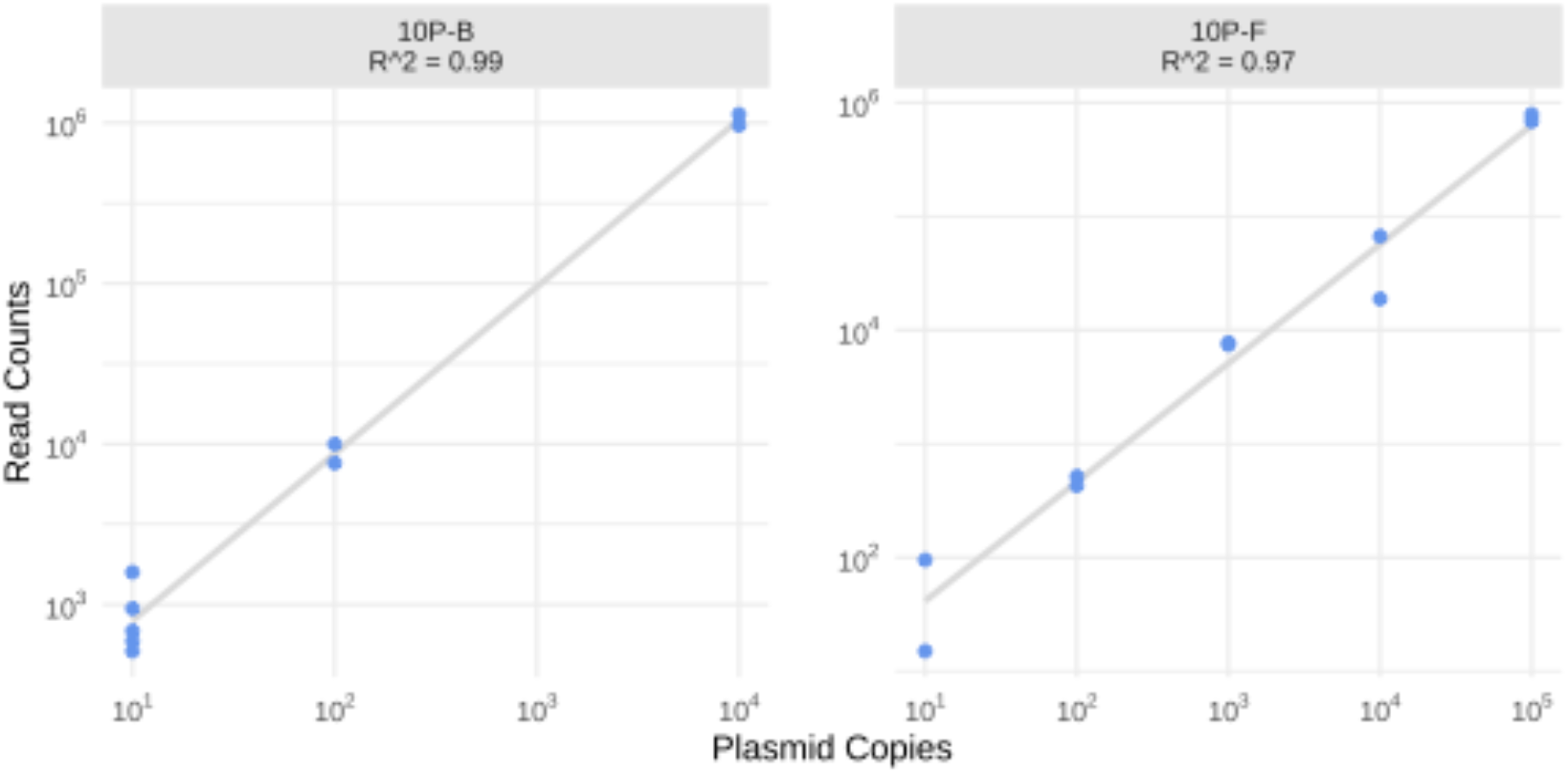
Linearity plots of 10-plasmid controls with L3 clustering parameter. The log10 transformed x-axis show the copy number of plasmid inputs, the log10 transformed y-axis represents L3 clustered read counts. Linear regression trendlines are plotted in gray, with corresponding R^2^ values. Linearity in 10-plasmid controls 10P-B and 10P-F is shown.

